# A Living Organoid Biobank of Crohn’s Disease Patients Reveals Molecular Subtypes for Personalized Therapeutics

**DOI:** 10.1101/2023.03.11.532245

**Authors:** Courtney Tindle, Gajanan D. Katkar, Ayden G. Fonseca, Sahar Taheri, Jasper Lee, Priti Maity, Ibrahim M. Sayed, Stella-Rita Ibeawuchi, Eleadah Vidales, Rama F. Pranadinata, Mackenzie Fuller, Dominik L. Stec, Mahitha Shree Anandachar, Kevin Perry, Helen N. Le, Jason Ear, Brigid S. Boland, William J. Sandborn, Debashis Sahoo, Soumita Das, Pradipta Ghosh

**Author notes:** **Senior corresponding authors: Brigid S. Boland, M.D.**; Associate Professor, Department of Medicine, University of California San Diego; 9452 Medical Center Drive, 1W 503, La Jolla, CA 92093. : **Email:** **William J. Sandborn, M.D.**; Professor, Departments of Medicine and Surgery, University of California San Diego; 9500 Gilman Drive, La Jolla, CA 92093. : **Email:** **Debashis Sahoo, Ph.D.**; Associate Professor, Department of Pediatrics, University of California San Diego; 9500 Gilman Drive, MC 0703, Leichtag Building 132; La Jolla, CA 92093-0703. : : **Email:** **Soumita Das, Ph.D.**; Associate Professor, Department of Pathology, University of California San Diego; 9500 Gilman Drive, George E. Palade Bldg, Rm 256; La Jolla, CA 92093. : **Email:**. **Senior corresponding and lead contact: Pradipta Ghosh, M.D.**; Professor, Departments of Medicine and Cell and Molecular Medicine, University of California San Diego; 9500 Gilman Drive (MC 0651), George E. Palade Bldg, Rm 232; La Jolla, CA 92093. : **Email:**. Equal contribution (Courtney Tindle, Gajanan D. Katkar, Ayden G. Fonseca, Sahar Taheri). Equal contribution (Soumita Das, Pradipta Ghosh). Department of Medical Microbiology and Immunology, Faculty of Medicine, Assiut University, Egypt (Ibrahim M. Sayed).

## Abstract

Crohn’s disease (CD) is a complex, clinically heterogeneous disease of multifactorial origin; there is no perfect pre-clinical model, little insight into the basis for such heterogeneity, and still no cure. To address these unmet needs, we sought to explore the translational potential of adult stem cell-derived organoids that not only retain their tissue identity, but also their genetic and epigenetic disease-driving traits. We prospectively created a biobank of CD patient-derived organoid cultures (PDOs) using biopsied tissues from colons of 34 consecutive subjects representing all clinical subtypes (Montreal Classification B1-B3 and perianal disease). PDOs were generated also from healthy subjects. Comparative gene expression analyses enabled benchmarking of PDOs as tools for modeling the colonic epithelium in active disease and revealed that despite the clinical heterogeneity there are two major molecular subtypes: immune-deficient infectious-CD [IDICD] and stress and senescence-induced fibrostenotic-CD [S2FCD]. The transcriptome, genome and phenome show a surprising degree of internal consistency within each molecular subtype. The spectrum of morphometric, phenotypic, and functional changes within the “living biobank” reveals distinct differences between the molecular subtypes. These insights enabled drug screens that reversed subtype-specific phenotypes, e.g., impaired microbial clearance in IDICD was reversed using agonists for nuclear receptors, and senescence in S2FCD was rectified using senotherapeutics, but not *vice versa*. Phenotyped-genotyped CD-PDOs may fill the gap between basic biology and patient trials by enabling pre-clinical Phase ‘0’ human trials for personalized therapeutics.

**GRAPHIC ABSTRACT:** 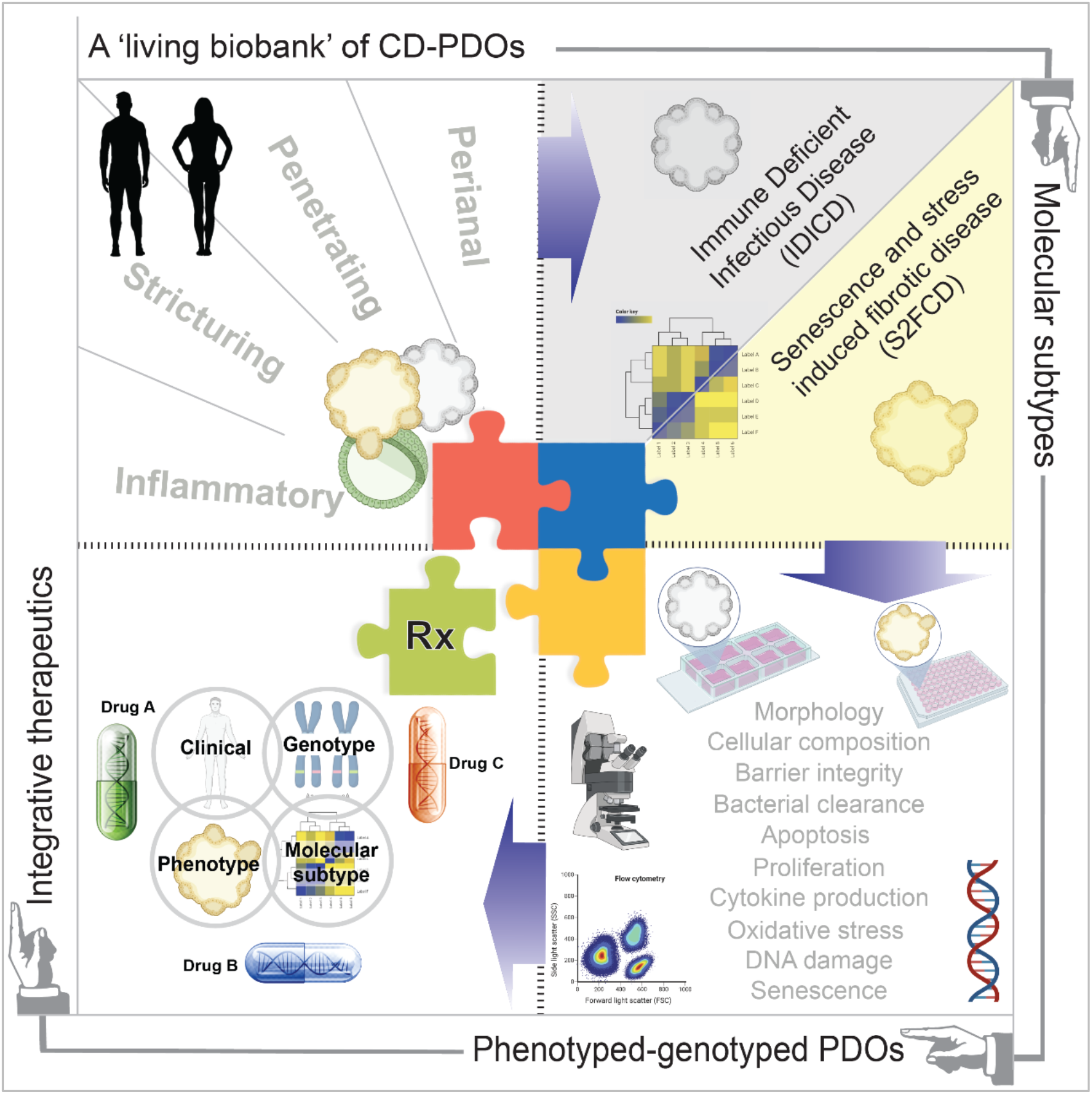

**In Brief:** This work creates a prospectively biobanked phenotyped-genotyped Crohn’s disease patient-derived organoids (CD-PDOs) as platforms for molecular subtyping of disease and for ushering personalized therapeutics.

**HIGHLIGHTS:**

- Prospectively biobanked CD-organoids recapitulate the disease epithelium in patients
- The phenome-transcriptome-genome of CD-organoids converge on two molecular subtypes
- One subtype shows impaired microbial clearance, another increased cellular senescence
- Phenotyped-genotyped PDOs are then used for integrative and personalized therapeutics

## INTRODUCTION

Crohn’s disease (CD) is a chronic incurable disease^1^. It is characterized by relentless progression with complications, e.g., intestinal fibrosis, penetrating fistulas and bowel destruction, that are fueled by inflammation^2^ and other malignant processes associated with such uncontrolled inflammation.

The unremitting inflammation in CD is believed to be multifactorial in origin^2^. Dysregulated interactions between the luminal microbiome, host genome, environmental triggers, and the gut immune system have been implicated. Although genetics (risk alleles/SNPs) can explain some variation in ileal vs ileo/colonic-predominant location of CD^3^, it has thus far failed to rationalize disease extent or behavior, response/resistance to therapeutics, predisposition to environmental triggers (infection, smoking) or risk of complications (e.g., extraintestinal manifestations and risk of colorectal cancers [CRCs])^3^. CD also lacks good preclinical animal models that faithfully recapitulate the diverse components of the human diseased tissue. Thus, heterogeneous behaviors of CD continue to pose a challenge and insights that can translate into personalization and precision in patient care are slow to emerge.

Recently, numerous studies have begun to unravel the role of different gut epithelial cell types in IBD using human patient-derived organoids (PDOs)^4, 5^. These studies have provided many new insights, e.g., the ability of PDOs to faithfully retain the tissue phenotypes in both CD ileum^6^ and in UC colon^7^, transcriptome and secretome^8^, the presence of stem cell dysfunction^9, 10^, telomere dysfunction^11^, increased apoptosis^12^, and impaired wound healing^13^ impaired barrier function^14, 15^. Almost all these studies suffer from the limited number of unique subjects studied, and they were either focused on UC or limited to the analyses of ileum of CD. None comprehensively studied the major clinical subtypes or disease behavior (Montreal classification B1-inflammatory, B2-stricturing and B3-penetrating, with/without perianal disease); thus, the molecular basis for such diverse clinical presentation has remained elusive. Consequently, most of the current FDA-approved therapeutic options^1^ lack personalization, and unsurprisingly, many randomized controlled trials (RCTs) fail to show statistically significant benefit over placebo in maintaining disease remission, especially in B3-penetrating/fistulizing disease^16^ (**Figure 1**; *Step 1*). Treatments with anti-inflammatory drugs may also be imprecise, i.e., they target proinflammatory cytokines without concrete evidence that excessive cytokines are the culprit while failing to tackle fundamental triggers (beyond inflammation) that initiate and perpetuate the disease.

**Figure 1.**
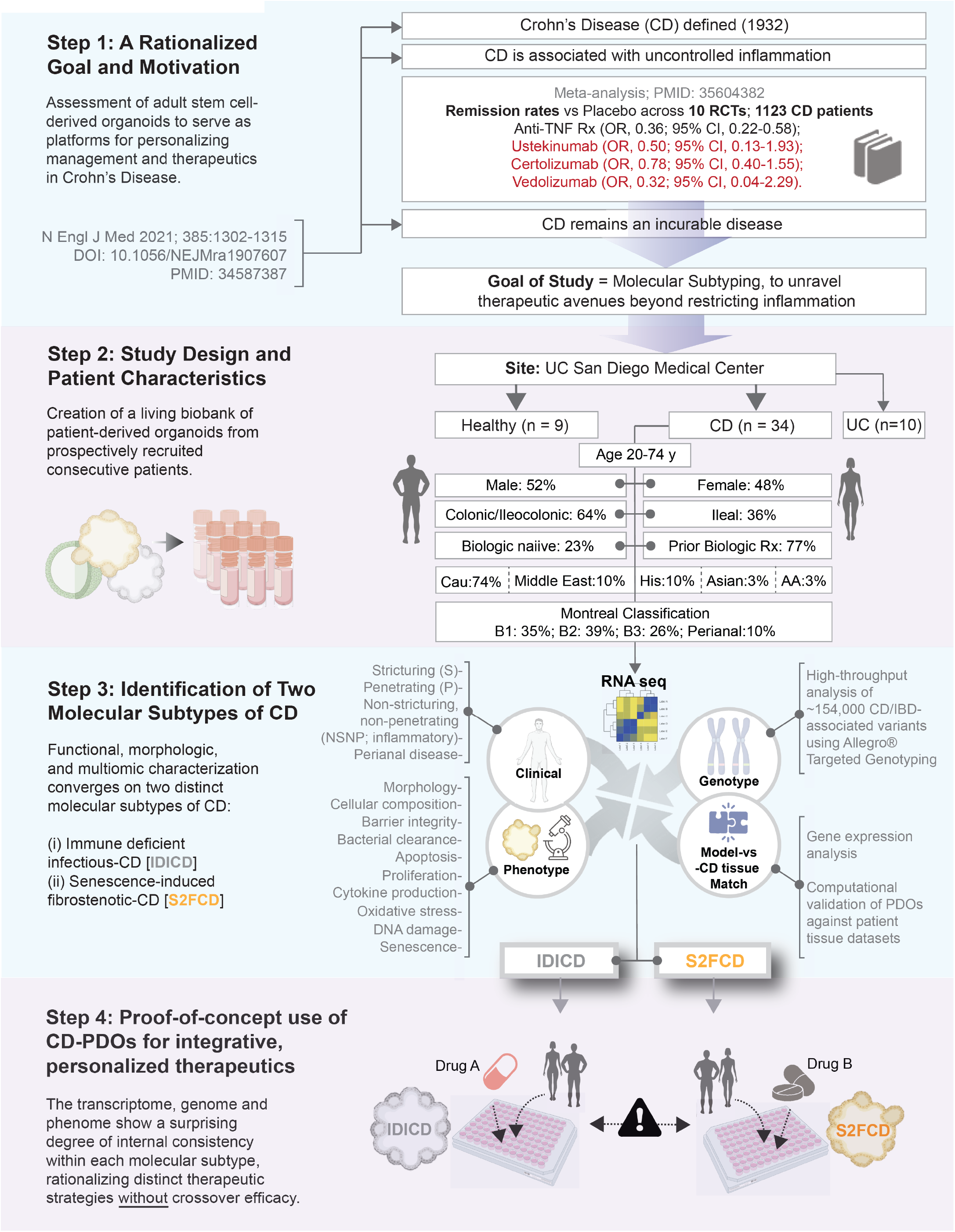
Study outline: Creation of a genotyped-phenotyped biobank of adult stem cell derived PDOs for enhancing personalized therapeutics in CD. Schematic presents key aspects of this study, starting with a rationalized goal and study motivation to tackle a complex disease with no good preclinical animal models that remains incurable (Step 1). Patients were enrolled in this study from the UC San Diego Health Center as source of colonic tissue biopsies for the isolation and creation of the CD-PDO biobank. Clinical, pathological and treatment history and the Montreal classification of disease were collected (see **Supplemental Information 1**). PDOs were generated from the adult stem cells at the crypt base, expanded and biobanked (in Step 2) for use in various assays (in Step 3). A total of n = 53 PDOs were used; 34 CD-PDOs from 34 unique CD subjects, 10 UC-PDOs (sampled for inflamed vs uninflamed locations of 4 subjects) and 9 heathy PDOs from 9 unique subjects. Various multiomic, morphologic and functional studies that were performed, some in low- and others in high-throughput modes (HTP; in 96-well plates) for systematic molecular and phenotypic characterization (catalogued in **Supplemental Information 2**). The study ends with proof-of-concept therapeutic studies (Step 4) in which drugs are rationalized and paired to each subtype with the intention to reverse in vitro the most prominent defect(s) within the subtype. Cross-over benefit of drugs across the subtypes is assessed and found as lacking. RCT, randomized controlled trials; OR, odds ratio; CI, confidence interval.

Here we sought to address some of these challenges by prospectively creating a living biobank of CD-PDOs generated from colon biopsies that is not just formidable in size [n = 53; 34 CD-PDOs, 10 UC-PDOs (sampled for inflamed vs uninflamed locations of 4 subjects) and 9 heathy PDOs], but also representative of the diverse patient population receiving care at any specialized tertiary care center in the United States (**Figure 1**; *Step 2*). Our approaches were geared to not just reveal new biology, but also sought to establish a way of ‘benchmarking’ CD-PDOs, through objective and stringent criteria, as reproducible and close ’replica’ of the diseased tissue (**Figure 1**; *Step 3*). Our findings highlight the potential of CD-PDOs to enable molecular subtyping of the disease and the use of phenotyped-genotyped CD-PDOs as platforms for pre-clinical Phase ‘0’ human trials for personalized and integrative therapeutics (**Figure 1**; *Step 4*).

## RESULTS AND DISCUSSION

### Establishment of a living CD biobank: study rationale and outline

Consecutive patients presenting to the UC San Diego Inflammatory Bowel Disease (IBD) Center were enrolled in this study. The only criteria were a clinically confirmed diagnosis of CD as the indication for an endoscopy and the ability to obtain informed consent. Detailed clinico-pathological and treatment-related information can be found in **Supplemental Information 1**. Patient characteristics are displayed in **Figure 1**; *Step 2*. All the major CD behaviors, as per the Montreal classification^17^, including the perianal modifier, were adequately represented (**Figure 1**; *Step 2*).

We focused on the colon because: (i) it is involved in ∼60% of patients with CD; while ∼half of those patients have synchronous involvement of the small intestine, the other half show disease that is limited to the colon^18^; (ii) most prior studies focused on the ileum and neglected the colon; none revealed disease subtypes.

To generate the biobank of PDOs, we adapted the culture conditions for long-term expansion of human colonic epithelium from *LGR5*-positive stem cells located at the bottom of the colon crypts along the lines of published work by others and us, with a few modifications (see *Methods*)^19–26^. Briefly, we modified the previously developed media^20–22^ that uses R-Spondin1 (the ligand for LGR5^27^), Noggin and Wnt (a ligand that is necessary for maintaining active crypt stem cells^28–30^) with minimal additives. All organoids could be readily expanded and frozen to create a living biobank with 100% success rate. Upon thawing, cell survival was typically > 90%, allowing us to systematically analyze them using ‘omics’, morphological, and functional studies, some of which were performed in low and several in high-throughput modes. Integration of these information with the clinical presentation led to the identification of two distinct molecular subtypes of CD (**Figure 1**; *Step 3*). Finally, PDOs were used in proof-of-concept studies as preclinical models to pair therapeutics with the intention to reverse key disease-driving cellular process(es) within each subtype, and to assess if subtype-paired therapeutics demonstrate cross-over efficacy across subtypes (**Figure 1**; *Step 4*).

### Objectivity in benchmarking PDOs as reproducible ‘replica’ of the diseased epithelium

Before using the PDOs in any functional studies, we analyzed them by bulk RNA seq to assess their ability to retain the altered gene expression pattern in the epithelium of the IBD-afflicted colon (**Figure 2A-D**; see **Supplemental Information 3**). Principal component analyses showed that while the healthy and CD-PDOs were distinct from each other, surprisingly, the CD-PDOs did not separate into 3 clinical subtypes. Instead, they segregated into two distinct clusters (CD1-gray and CD2-yellow; (see Factor map, **Figure 2B**). All the penetrating (P, B3) CD-PDOs and PDOs from perianal disease were in the gray cluster (**Supplemental Information 3**), however, the NSNP (B1) and stricturing (B2) CD-PDOs were split between the gray and the yellow clusters (**Figure 2B**). All the ten ulcerative colitis (UC)-PDOs that were simultaneously isolated and analyzed by RNA seq were found to co-cluster with CD-PDOs in the gray cluster (**Supplementary Figure 1A-B**), regardless of whether they were derived from the involved or uninvolved segments of the colon. Results indicate that UC may have shared pathophysiology with one of the two CD subtypes, but both are distinct from the healthy PDOs.

**Figure 2.**
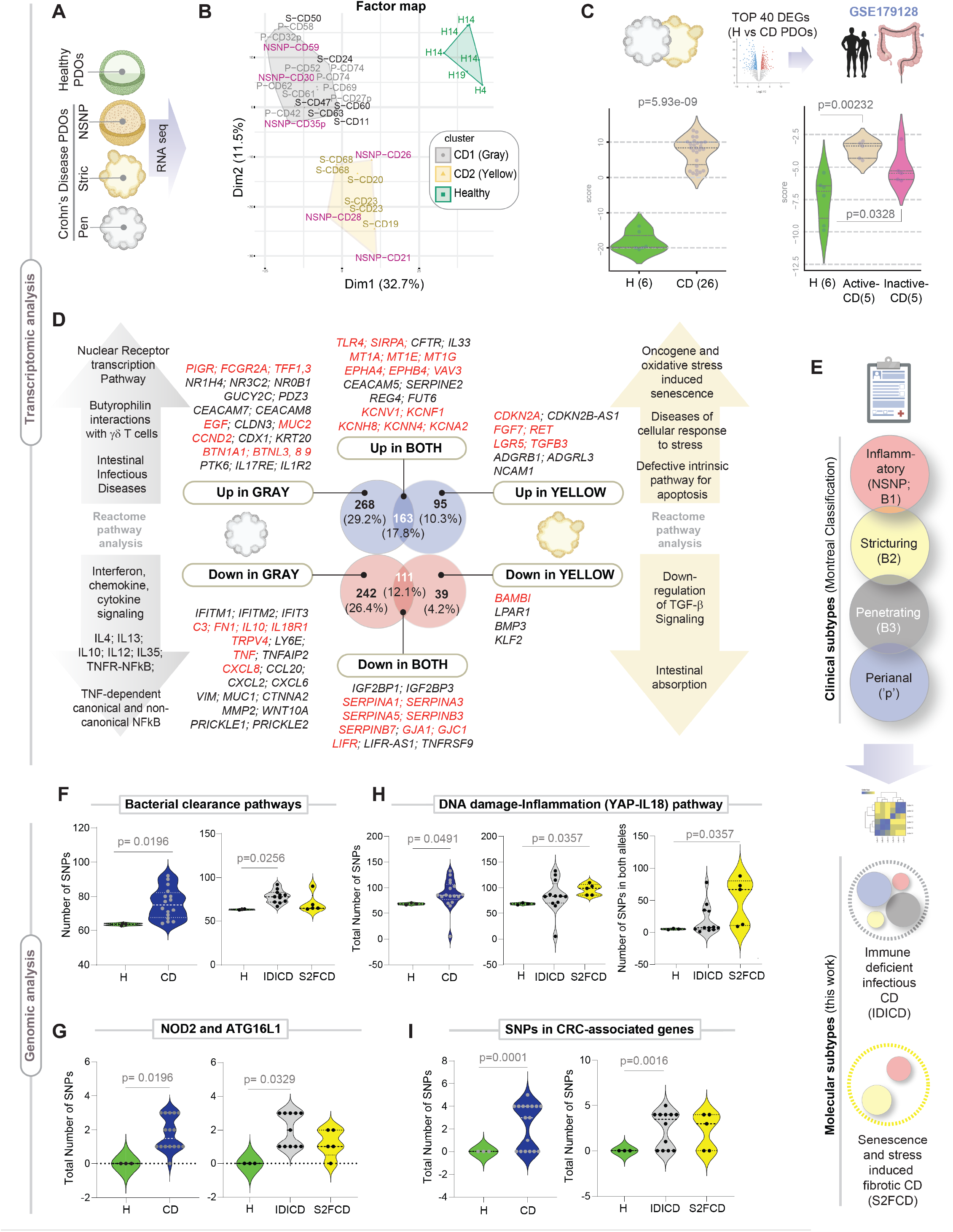
Transcriptome and genome analyses of CD-PDOs reveal the existence of two distinct molecular subtypes of CD. **A.** Schematic showing the overall rationale and study design for the transcriptomic studies on healthy and CD- PDOs. **B.** A factorial map generated by performing the Hierarchical Clustering on principal components (HCPC) analysis is plotted onto the first two dimensions. HCPC was used to compute hierarchical clustering on principal components indicates the division of the CD-PDOs into two distinct clusters: gray and yellow, which are differentiated from healthy controls (green). Individual samples are labeled in the factor map (patient information for the corresponding samples are in (see **Supplemental Information 3**). H, healthy; NSNP, non-stricturing, non-penetrating; S, stricturing; P, penetrating. Samples (annotated with ‘p’) were from subjects with perianal disease. See also **Supplementary Figure 1** for comparison with UC-PDOs. **C.** *Top*: Schematic outlines the strategy used to objectively assess the ability of CD-PDOs to recapitulate the microdissected epithelium from the colons of CD patients (GSE179128). *Bottom*: Violin plots show the composite score of the PDO-derived top upregulated DEGs (left) and in the laser capture microdissected colonic epithelium (right). Values in parenthesis indicate unique patients. See **Supplemental Information 4** for list of DEGs. **D.** Differentially expressed (UP or DOWN) genes uniquely in the gray (CD1; *left*) vs yellow (CD2; *right*) cluster of CD-PDOs, or those that are shared between both subtypes (CD1+2) are listed, alongside the enriched pathways they represent. See **Supplemental Information 4** for a complete list of DEGs and reactome pathways. See also **Supplementary Figure 2-3** for additional PCOA and reactome pathway enrichment analyses and **Supplementary Figure 4-5** for transcriptome-derived insights into CD-associated changes in the crypt-axis differentiation score, stem-cell dysfunction, cellular composition, and other properties. **E.** Schematic summarizes how various clinical subtypes of CD (Montreal classification) fit into two broad molecular subtypes, immune-deficient infectious CD and senescence and stress-induced fibrotic CD. **F-I**. Violin plots show the number of mutations in genes within the indicated pathways (see **Supplemental Information 5** for gene list) in CD-PDOs vs healthy controls. Plots on the left compare all CD-PDOs combined, whereas plots on the right separate the CD-PDOs by molecular subtypes, IDICD and S2FCD. Plots in G specifically display the frequency of NOD2 SNPs rs2066843 and rs2076756 and ATG16L1 SNP rs2241880 in CD-PDOs vs healthy controls. Statistical significance was assessed by Mann-Whitney (F, G-*left*, H, I), one way ANOVA (G-*right*, I-*right*), Welch’s test (I). Only significant *p* values are displayed. See also **Supplementary Figure 6**.

An analysis of differentially expressed genes (DEGs) revealed the common set of genes and cellular pathways and processes that are up- or down-regulated in both gray and yellow clusters (**Supplemental Information 4** for the list of DEGs and pathways). To objectively assess how well PDOs reproduced the altered epithelial biology in the setting of active disease, we took advantage of a high-quality publicly available dataset of laser microdissected IBD-afflicted colonic epithelia (GSE179128)^31^; as one of a kind, this dataset included both active and inactive states of UC and CD. The top upregulated DEGs in CD-PDOs were induced in the micro-dissected CD- (**Figure 2C**) and UC- (**Supplementary Figure 1C**) afflicted colonic epithelia, exclusively in the active disease state. Similarly, the top upregulated DEGs in UC-PDOs were induced in the microdissected IBD-afflicted colonic epithelia exclusively in the active disease state (**Supplementary Figure 1D**). These unbiased assessments of the ‘match’ between gene expression patterns in PDO vs tissue indicated that the CD/UC-PDOs replicate the disease activity in the colonic epithelium of patients. When IBD organoids from other groups were analyzed using the same unbiased yardstick on the same tissue cohort, some reproduced the disease state^32, 33^ (**Supplementary Figure 1E**), but others did not (**Supplementary Figure 1F-G**). Reproducibility was seen in studies where adult stem cells from the colon crypt were used to derive the PDOs and L-WRN conditioned media was used for their expansion and biobanking. Reproducibility was lacking in PDOs grown in defined media^11, 24^ or where PDOs were derived from induced pluripotent stem (iPS) cells^34^.

### Identification of two distinct molecular subtypes of CD

Next, we compared the gene expression patterns in the gray vs. yellow clusters (**Figure 2D**) and asked what cellular pathways or processes may be enriched in the genes that are uniquely up- or down-regulated in each subtype (see **Supplemental Information 4** for a complete list of genes and reactome pathways; **Supplementary Figure 2-3** for PCA, heatmap and reactome analyses). Uniquely upregulated in the gray cluster were genes that are involved in intestinal infectious diseases and the butyrophilins; the latter are molecules that are used by epithelial cells to shape organ-specific γδ T cells^35^ (**Figure 2D**-*top left*). Uniquely downregulated in the gray cluster were genes that are involved in all the major interferon, chemokine and cytokine signaling pathways (**Figure 2D***-bottom left*). As for the yellow cluster, the uniquely upregulated genes were, among others, *CDKN2A*, *LGR5* and *RET,* all known to be involved in oncogene and oxidative stress-induced senescence, cellular response to stress and defects in apoptosis (**Figure 2D***-top right*). Uniquely downregulated in the yellow cluster were genes such as *BAMBI*, the potent endogenous inhibitor of fibrogenic TGFβ signals^36^ ( **Figure 2D***-bottom right*). The up-regulated DEGs shared between the two subtypes were notable for the pattern recognition receptor, *TLR4* and its negative regulator and the self-recognition molecule, Signal Regulatory Protein Alpha (*SIRPA*; a.k.a *CD47*), metallothioneines, *MT1A, MT1E, MT1G*, the Ephrins *EPHA4*, *EPHB4*, voltage gated potassium channels, and the RhoGTPase and oncogene, *VAV3* (**Figure 2D**). The downregulated DEGs shared between both subtypes were notable for numerous serine protease inhibitor genes (*SERPINEA1, -A3, -A5; -B3, -B7*) and gap junction genes *GJA1* and *GJC1*. We also observed downregulation of *TNFSF9* (aka, CD137), which is known to enhance epithelial TJ resistance^37^.

These results indicate that even though CD has 4 major clinical behaviors as per Montreal classification (**Figure 2E***-top*), these heterogeneous presentations fit into one of two molecular subtypes (**Figure 2E***-bottom*). Because the gray cluster uniquely represented an infectious disease-like state in the setting of reduced cytokine/inflammatory responses (i.e, paradoxical immune deficiency), we named this subtype immune-deficient infectious CD (IDICD). Because the yellow cluster uniquely represented stress and senescence in the setting of reduced anti-fibrogenic signaling, we named this subtype senescence and stress induced fibrotic CD (S2FCD).

The transcriptome analyses also revealed a plethora of other clues into the shared and distinct epithelium-intrinsic defects in the two molecular subtypes (detailed in **Supplementary Text**). Briefly, CD-PDOs showed skewed crypt differentiation, more prominently in IDICD (**Supplementary Figure 4A-E**), stem cell dysfunction (**Supplementary Figure 4F-G**), lower paneth:goblet cell ratio and goblet cell dysfunction (**Supplementary Figure 5A-C**), lower expression of TJ- (**Supplementary Figure 5D**), and impaired expression of DNA damage response (**Supplementary Figure 5E**) and mitotic checkpoint (**Supplementary Figure 5F**) genes. A comprehensive analysis also revealed subtype-specific dysregulation of the expression of genes nearest to every CD-risk variant significantly associated with CD based on GWAS (see *Methods*; **Supplementary Figure 5J**).

### The transcriptome and genome converge within each molecular subtype of CD

Genetic contributions to CD behavior remain poorly defined to date^3^ (summarized in **Supplementary Figure 6A**). We isolated genomic DNA from healthy and CD-PDOs for targeted sequencing of ∼154,000 SNPs that have been implicated previously in IBD (see Methods; **Supplementary Figure 6B**). SNPs in genes within the bacterial clearance pathway (see **Supplemental Information 5**) were over-represented in the CD-PDOs, specifically in the IDICD subtype (**Figure 2F**). More specifically, CD-risk alleles in *NOD2* and *ATG16L1* were over-represented in the IDICD CD-PDOs (**Figure 2G**). By contrast, SNPs in genes within a senescence and DNA-damage-induced *YAP-IL18* proinflammatory pathway^38^ were over-represented in the S2FCD PDOs (**Figure 2H**). CRC-associated mutations were over-represented in the CD-PDOs, primarily in the IDICDs (**Figure 2I**); these results agree with epidemiologic studies showing higher CRC risk in penetrating (B3)^39, 40^ and perianal fistulizing CD^41^. A higher frequency of SNPs were also found in monogenic risk alleles-- *RET* (**Supplementary Figure 6C**) and *POU5F1* (**Supplementary Figure 6D**); both genes have been associated with early onset CD^42, 43^. *RET* supports cell-cell adhesion to resist TNFα-challenge^44^, and mutations in this gene has been implicated in the development of enterocolitis independent of NOD2^45^. Although, in most of the other pathways analyzed, CD-PDOs harbored more mutations, we did not find a subtype specific enrichment, i.e., apoptosis (**Supplementary Figure 6E**), DNA damage (**Supplementary Figure 6F**), proliferation (**Supplementary Figure 6G**) and epithelial TJs (**Supplementary Figure 6H**).

Because the SNPs that impair bacterial clearance are enriched in the IDICD subtype, and the SNPs within the senescence associated DNA damage and inflammation are enriched in the S2FCD subtype (see **Table 1**), we conclude that the transcriptome and genome converge on related themes within each molecular subtype.

**Table 1.** A summary of multi-omics, functional and cell biological approaches used, and major statistically significant findings reported in this work. NT, not tested; HTP, high throughput; LTP, low throughput

| Molecular Subtype | IDICD<br>(Immune-deficient Infectious CD) |  |  |  | S2FCD<br>(Senescence and stress induced fibrotic CD) |  | Approach | Comments |
| --- | --- | --- | --- | --- | --- | --- | --- | --- |
| **Montreal Classification** | Penetrating (all) | NSNP (some) | Perianal (all) | Strictureing (few) | Strictureing (most) | NSNP (some) | RNA seq analysis of CD-PDOs vs Healthy | n/a |
| **Barrier integrity** | Intact | Impaired | Intact | Intact in most, impaired in some | Intact in most, impaired in some | Impaired | TEER measurement *<br>FITC-Dextran leakage * | * = HTP in 96-wells |
| **Dysmorphic growth** | no | Yes; defect in lumen formation | no | Yes; asymmetry, high cellularity | Yes; asymmetry, high cellularity | yes | Light microscopy*, IMARIS imaging* | * = HTP in 96-wells |
| **Apoptosis** | High | normal | NT | High (easily seen by H&E) | High | Normal | TUNEL assay# | # = LTP-semi-HTP in 8-wells |
| **Apoptosis in response to TNFα** | High | High | NT | High | High | High |  |  |
| **Proliferation** | High | normal | NT | High | High | normal | BrdU incorporation * | * = HTP in 96-wells |
| **Ox DNA/RNA damage** | normal | High | NT | High | High | High | ELISA based assay * | * = HTP in 96-wells |
| **Crypt cell composition, differentiation** | High crypt-axis score, high CEACAM7+ brush border cells, high goblet cells; defective MHC-II+ ISC-III, high goblet cells |  |  |  | Low crypt-axis score, defective MHC-II+ ISC-III, high goblet cells; defect in mitotic checkpoint. |  | RNA seq | n/a |
| **Paneth cell dysfunction** | Yes (excessive degranulation) |  |  |  | Yes (excessive degranulation) |  | Imaging (IF and EM) | LTP |
| **Goblet cell AMP production** | Impaired severely |  |  |  | Reduced slightly |  | RNA seq | n/a |
| **DNA damage (dsDNA break)** | Normal |  |  |  | High |  | Flow cytometry (gH2AX) | LTP |
| **Senescence** | Normal | | | | High | | SPIDER β-Gal assay, Flow cytometry | # = LTP-semi-HTP in 8-wells |
| **Bacterial clearance** | Impaired |  |  |  | NT |  | Infection, lysis, plating and colony counting | LTP on EDMs |
| **Induction of ROS** | High |  |  |  | Very high |  | ELISA * | * = HTP in 96-wells |
| **Microbe-challenged production of cytokines** | Impaired |  |  |  | Normal |  | Multiplex ELISA assays [Mesoscale discovery] * | * = HTP in 96-wells |
| **Genotype-phenotype relationship** | SNPs in genes within the bacterial clearance pathway, NOD2/ATG16L1 disease-driving SNPs, and SNPs in CRC-associated genes. |  |  |  | SNPs in the genes within the DNA damage-YAP-IL18 inflammatory pathway. |  | Genotyping * and SNP analysis | HTP |

NT, not tested; HTP, high throughput; LTP, low throughput

### The molecular subtypes of CD-PDOs display shared and distinct phenomes

We next asked if the convergent transcriptome and genome support a cohesive phenome in the CD-PDOs; such cohesion, if present, is expected to offer two distinct advantages. First, convergence of genome-transcriptome- phenome could help rationalize and prioritize therapeutic agents personalized to the molecular subtype of the CD-PDOs. Second, the rescue of the phenome in response to an intervention may serve as an objective metric of therapeutic response. We tested this concept using the two molecular subtypes of CD-PDOs.

#### S2FCD PDOs show dysmorphic growth features-

Unlike healthy colonoids, CD-PDOs presented with a range of patient-specific morphologies in 3D growth conditions, ranging from thin-walled cystic structures to complex structures with thick walls and off-centered lumen or multiple lumens, to compact organoids with either smaller lumen or completely devoid of a lumen (**Figure 3A**; **Supplementary Figure 7A**). Observations by light microscopy revealed that CD-PDOs were less likely to grow as thin-walled organoids with central lumen, and more likely to grow into solid structures (no lumen) compared to their healthy counterparts (**Supplementary Figure 7B***-left*). S2FCD showed the most dysmorphic growth, whereas IDICD PDOs were the closest to healthy PDOs (**Figure 3B**). A clinical subgroup analysis showed that both NSNP (B1)- and stricturing (S, B2) CD-PDOs were dysmorphic and the penetrating (P, B3) CD-PDOs were the least (**Supplementary Figure 7B***-right*). Findings by light microscopy were verified by quantitative fluorescence microscopy (**Supplementary Figure 7C**). NSNP (B1)- and stricturing (S, B2) CD- PDOs consistently showed the most volume in 3D (voxel count, **Supplementary Figure 7D**), higher cellularity (nuclear count; **Supplementary Figure 7E**) and asymmetry (Bounding box analyses; **Supplementary Figure 7F-G**), whereas ellipticity and sphericity were comparable across all healthy and CD-PDOs (**Supplementary Figure 7H-I**). H&E staining also revealed the presence of frequent apoptotic nuclear events, however, was unique to the stricturing (S, B2) CD-PDOs (**Figure 3A***, arrows*). These analyses show that adult stem cell derived organoids display abnormal 3D growth.

**Figure 3.**
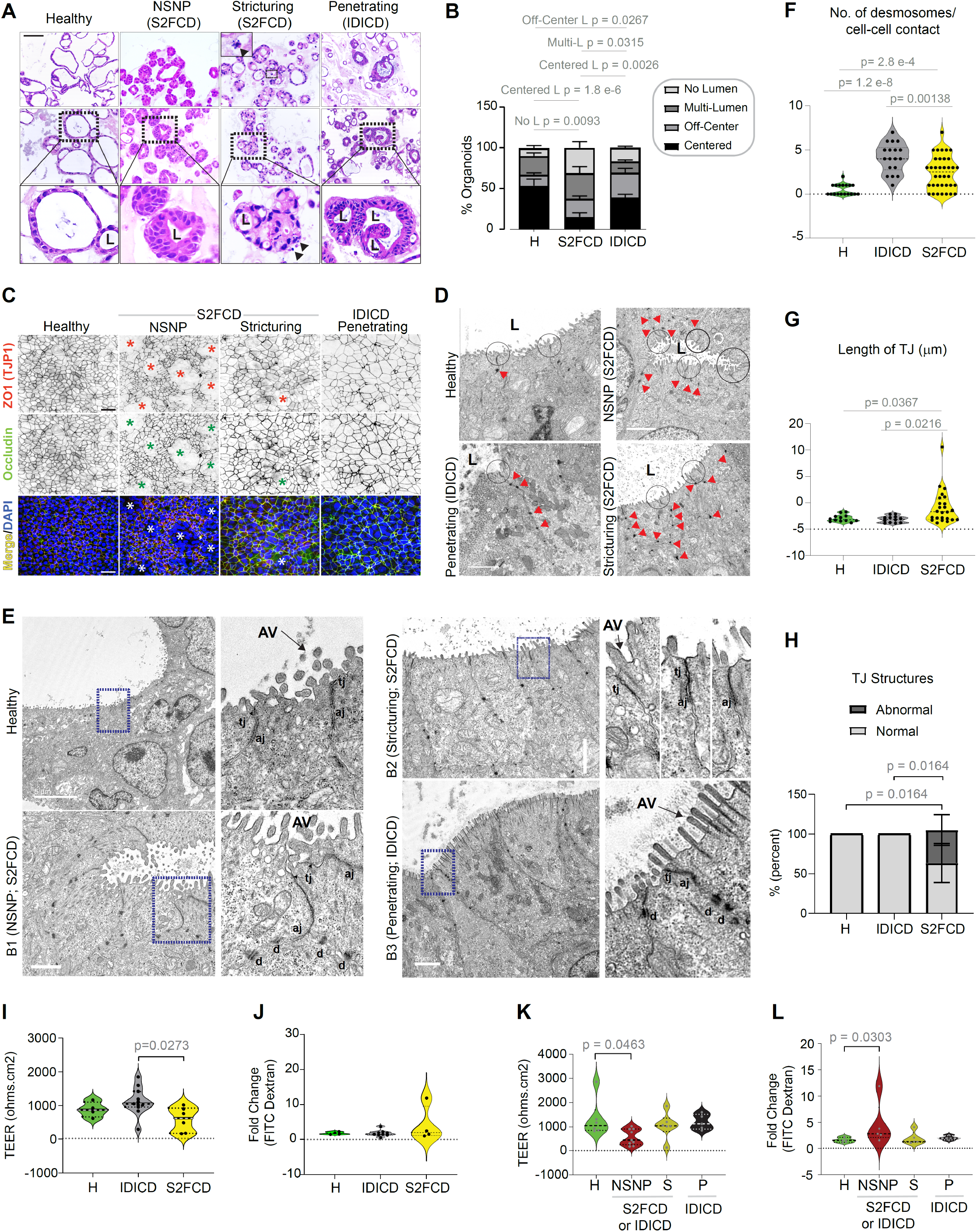
Assessment of morphology and barrier integrity of CD-PDOs. **A-B.** Representative images (A) of hematoxylin and eosin-stained FFPE-CD-PDOs are shown from each clinical subtype of CD alongside healthy controls. Scale bar = 100 µm. Top and middle rows show two examples from each clinical subtype, whereas bottom panel shows a magnified region from middle panel. L = lumen. Arrowhead = nuclear fragmentation (likely apoptotic bodies). Stacked bar plots in B shows the quantification of the proportion of each type of organoid structure in various CD subtypes (See **Supplementary Figure 7B** for all CD subtypes combined; B-*right,* separated into CD subtypes). Statistical significance was assessed by one way ANOVA. Only significant *p* values are displayed (n = 3-8 in each group). See **Supplementary Figure 7** for morphologic assessment across all clinical CD subtypes, which includes quantitative morphometrics using IMARIS and lumen position/presence by light microscopy. **C.** Various clinical subtypes of CD-PDOs were differentiated into polarized monolayers on transwells (enteroid- derived monolayers; EDMs), fixed and stained for ZO1 (red) and occludin (green) and DAPI (nuclei, blue) and analyzed by confocal microscopy. Representative fields from different CD subtypes are shown; individual red and green channels are displayed in grayscale. Asterisk = areas of impaired barrier. Scale bar = 50 µm. **D.** Electron micrographs of healthy and CD-PDOs display apical cell-cell junctions. Red arrowheads = desmosomes. Scale bar = 2 µm. **E.** Electron micrographs of tight junctions in healthy and CD-PDOs of various subtypes are shown. Boxed region on the left is magnified on the right. AV, apical villi; tj, tight junction; aj, adherens junction; d, desmosomes. Scale bars = 5 µm (top two panels) and 2 µm (bottom two panels). **F-H**. Plots display the quantification of no. of desmosomes/cell-cell contact (F), the length of TJ (G) and the frequency of abnormal defects/TJ structure (H) observed by TEM. Statistical significance was assessed by one way ANOVA (n = 7-13 fields analyzed in each subtype of PDO). See **Supplementary Figure 8A-C** for the same analysis displayed as clinical subtypes of CD. **I**-**L**. Violin plots show the fold change in TEER across (I, K) and FITC dextran leakage through (J, L) CD-EDMs compared to healthy controls. Data are displayed either as molecular (I, J) or clinical (K, L) subtypes. Statistical significance was assessed by one way ANOVA. Only significant *p* values are displayed (n = 5-9 subjects in each group, 2-5 repeats in each PDO). See **Supplementary Figure 8D-F** for all CD subtypes combined. See **Supplemental Information 2** for subjects analyzed in each assay.

#### Inflammatory CD-PDOs in the S2FCD molecular subtype display barrier dysfunction-

We prepared enteroid-derived monolayers (EDMs) from healthy and CD-PDOs using established methodologies^22, 46, 47^ that were previously successfully adapted for PDOs from patients with inflammatory bowel disease^20, 48^. Once established as stable monolayers (i.e., trans-epithelial electrical resistance [TEER] reaches plateau over 3 consecutive readings), we analyzed the EDMs for tight junctions (TJs) by confocal microscopy (**Figure 3C**) and TEM (**Figure 3D-H**) and for paracellular permeability using two well-established functional assays, leakage of fluorescently labeled dextran^49^ and trans-epithelial electrical resistance^50^ (**Figure 3I-K**). The TJ proteins, ZO1 and occludin were readily visualized in EDMs from all CD subtypes; while 80% of NSNP (B1)- and ∼25-30% of stricturing (S) CD-EDMs showed breaks in the monolayer (asterisk, **Figure 3C**), none of the penetrating (IDICD) EDMs showed the same. TEMs revealed increased number of desmosomes in both IDICD and S2FCD subtypes (**Figure 3D-E**). The TJs were short, and the membrane leaflets of the neighboring cells were tightly opposed in healthy PDOs (**Figure 3D**); however, they were either elongated or showed interrupted membrane opposition in the S2FCD PDOs (**Figure 3C-D**). As for the IDICD PDOs, the TJs and AJs appeared morphologically indistinguishable from healthy PDOs, with two notable exceptions: (i) desmosomes were increased and (ii) apical microvilli were elongated and prominent in brush border cells (**Figure 3D**). Quantification of these features confirmed that although these aberrant junctions were more frequently encountered in the S2FCD PDOs (**Figure 3E-G**), the NSNP-subtype were the primary contributors to the observed differences (**Supplementary Figure 8A-C**).

Paracellular permeability was not increased in CD-EDMs compared with healthy-EDMs (**Supplementary Figure 8D-F**). The S2FCD-PDOs appeared to form a slightly leakier monolayer than IDICD-PDOs, apparent only by TEER assessment (**Figure 3I-J**). A clinical subtype analysis, however, confirmed that NSNP (B1)-EDMs are significantly leakier than healthy EDMs based on both lower TEER assessments (**Figure 3K**) and the recovery of higher amounts of FITC-dextran from the basolateral side (**Figure 3L**). These results indicate that in the absence of additional stressors, barrier integrity in the CD-colon may be largely comparable to that seen in the healthy colon, except for the inflammatory (NSNP/B1) CD-subtype.

#### IDICD harbors Paneth cell defects, whereas S2FCD shows oxidative stress and DNA damage-

We next assessed the CD-PDOs for epithelial indicators of inflammation and stress, i.e., altered mucin production and cell composition^51^ and genotoxic stress^52^. *MUC2* transcript, a marker of goblet cells, was elevated in CD-PDOs, primarily in the IDICD PDOs, as determined by qPCR (**Supplementary Figure 9A**; **Figure 4A**). Lysozyme (*LYZ*), a marker of the Paneth cells, was simultaneously assessed in the same PDOs; besides their antimicrobial function, these cells are essential for the growth of LGR5+ stem cells^53, 54^ and abnormal production of lysozyme is known to regulate the inflammatory milieu in colitis^55^. Although the transcripts of lysozyme were unchanged in CD-PDOs (**Supplementary Figure 9B**), we observed a reduced *LYZ*:*MUC2* ratio, indicative of a skewed ratios of secretory cells-- Paneth and goblet cell—in CD-PDOs, largely attributable to IDICD PDOs (**Figure 4B**). Confocal immunofluorescence studies showed reduced lysozyme-positive cells but an increased presence of luminal lysozyme and mucin-producing goblet cells in all CD-PDOs (**Figure 4C**). Enhanced degranulation and granule depletion was confirmed at a higher resolution by transmission electron microscopy (TEM; **Figure 4D**). Because interferon (IFN)γ induces Paneth cell degranulation^53, 56^, and because high serum IFNγ levels in CD originate from the immune cell infiltrates in the intestine^57^, it was surprising that our CD-PDOs retain this phenotype in culture despite being removed from those cells. Regardless, the findings agree with prior reports of enhanced extrusion and Paneth cell degranulation^53, 56^ increased thickness of mucin^58^ and the number of goblet cells^59^ in CD. Significant expansion of enteroendocrine cells was found in all CD-PDOs across every clinical subtype (**Supplementary Figure 9C**; **Figure 4A**), as determined by the abundance of *CHGA* transcripts, which is consistent with prior reports of observed increases in CD tissues^60^. Sucrase isomaltase (*SI*), which is a marker of the brush border enterocyte^61^ is increased in CD-PDOs, specifically in the penetrating (P, B3) CD- PDOs (**Supplementary Figure 9C**; **Figure 4A**).

**Figure 4.**
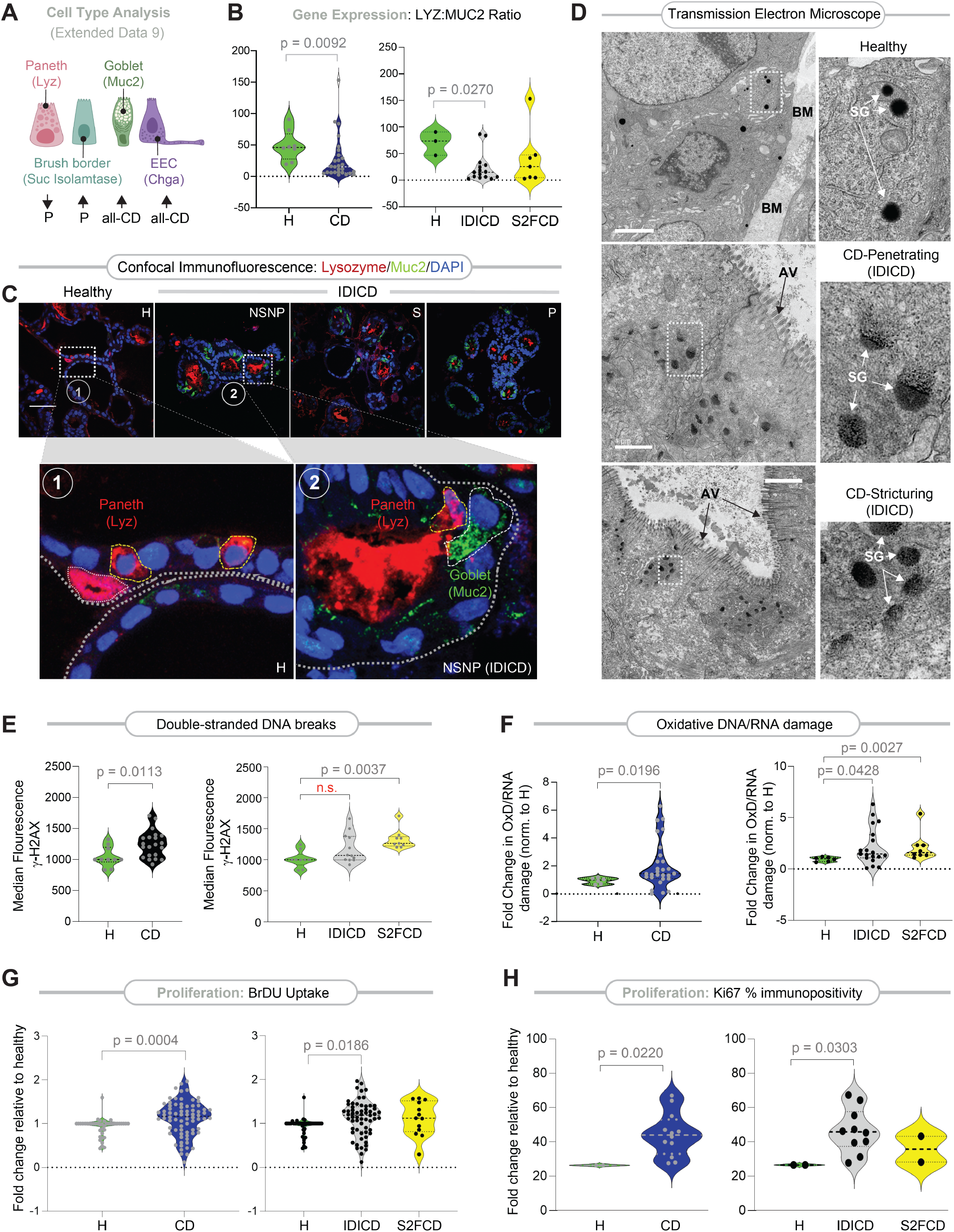
CD-PDOs retain evidence of altered cell composition, high oxidative stress, and turnover. **A.** Schematic summarizing the relative expression of type, as determine by gene expression (markers used for each cell type in parenthesis). Up-arrow = upregulation. Down-arrow = downregulation. P = penetrating-CD. See **Supplementary Figure 9A-D** for violin plots displayed as both clinical and molecular subtypes of CD. **B.** Violin plots show the ratio of LYZ and MUC2 transcripts in CD-PDOs vs healthy controls [B-*left,* all CD subtypes combined; B-*right,* separated into molecular subtypes of CD]. Statistical significance was assessed by Mann Whitney. Only significant *p* values are displayed (n = 6-15 subjects in each group). See **Supplementary Figure 9E** for the same data, displayed as clinical subtypes of CD. **C.** FFPE of CD-PDOs of the IDICD subtypes were analyzed for goblet (MUC2; green) and Paneth (Lysozyme; red) cells by confocal immunofluorescence. Representative images are shown. Scale bar = 150 µm. Boxed regions (numbered 1, 2 in upper panels) are magnified below. See **Supplementary Figure 9F** for quantification of images and **Supplementary Figure 9G** for summary. **D.** Electron micrographs of Paneth cells in healthy and CD-PDOs of IDICD subtypes are shown. Boxed region on the left is magnified on the right. BM = basement membranes; AV, apical villi; SG, secretory granules. Scale bar = 1 µm in top and middle panels, and 2 µm in the bottom panel. **E.** Violin plots show the extent of DNA damage, as determined by flow cytometry analysis of γH2AX in CD-PDOs vs healthy controls [E-*left,* all CD subtypes combined; E-*right,* separated into molecular subtypes of CD]. Statistical significance was assessed by Mann Whitney (left) or one way ANOVA (right). Only significant *p* values are displayed. **F.** Violin plots show the extent of oxidative DNA/RNA damage, as determined by ELISA on the supernatants from CD-PDOs vs healthy controls. Statistical significance was assessed by Mann Whitney. Only significant *p* values are displayed. See **Supplementary Figure 10A-B** for the display of the findings based on clinical subtypes of CD. **G.** Violin plots show the extent of BrDU incorporation over 24 h on four-day old CD-PDOs grown in 96-well plates prior to assessment by ELISA. Statistical significance was assessed by Mann-Whitney (left) and one way ANOVA (right). Only significant *p* values are displayed (n = 5-8 subjects in each group). See **Supplementary Figure 10C-D** for the display of the findings based on clinical subtypes of CD. **H.** Violin plots show % cells with Ki67-positive nuclei in CD-PDOs vs healthy controls, as determined by particle analysis on confocal images of Ki67-stained PDOs. Statistical significance was assessed by Mann-Whitney. Only significant *p* values are displayed (n = 2-5 subjects in each group). See **Supplementary Figure 10E-F** for the display of the findings based on clinical subtypes of CD. See **Supplemental Information 2** for subjects analyzed in each assay. See also **Supplementary Figure 11** for cellular apoptosis in CD-PDOs at baseline and upon challenge with TNFα.

Compared to healthy PDOs, CD-PDOs showed increased genotoxic stress. Increases were seen in both double-stranded DNA breaks (**Figure 4E-***left*) and oxidative nucleotide damage (**Figure 4F-***left*), as determined by γH2AX-based flow cytometry and in ELISA-based assays that measure the end products of oxidative RNA/DNA damage, respectively. S2FCD PDOs largely contributed to this increase (**Figure 4E-***right***, F-***right*). As for clinical subtypes, increased genotoxic stress was primarily encountered in NSNP (B1)- and more prominently in stricturing (S, B2) CD-PDOs, whereas penetrating CD were relatively spared (**Supplementary Figure 10A- B**).

That CD-PDOs retain DNA damage, ‘Paneth cell degranulation’ and altered cellular composition phenotypes *ex vivo* suggests the retention of some epigenetic memory in the adult stem cell, likely imprinted by IFNγ, tumor necrosis factor (TNF)α, or other stressful triggers from the diseased tissue of origin.

#### IDICD shows higher cell proliferation and turnover -

Because barrier loss due to increased death and/or impaired regeneration is a constant feature in inflammatory colitis^62^, we next analyzed these properties in CD-PDOs. Two complementary approaches were used— estimation of Bromodeoxyuridine (5-bromo-2’-deoxyuridine (BrdU) incorporation in colorimetric assays (**Supplementary Figure 10C**) and by staining FFPE organoids for the Ki67 protein (MKI67; **Supplementary Figure 10E**), a reliable marker of proliferation in the colon crypts^63^. Both assays agreed, in that, CD-PDOs display higher proliferation (**Figure 4G-***left,* **4H-***left*); however, a subtype-specific analysis showed that increased proliferation was primarily and most consistently encountered in the IDICD subtype (**Figure 4G-***right***, 4H-***right*). Among the clinical subtypes in IDICD, stricturing (S, B2) CD-PDOs showed the most proliferation (**Supplementary Figure 10D, 10F**).

When the same PDOs under similar conditions as above were assessed for apoptosis by TUNEL assays (**Supplementary Figure 11A**), apoptosis was found to be increased at baseline in CD-PDOs (all subtypes combined); **Supplementary Figure 11B***-left*). Subtype-specific analysis showed that apoptosis is more pronounced in the stricturing- (S, B2) and penetrating- (P, B3) subgroups (**Supplementary Figure 11B***-right*), but similar across the molecular subtypes (**Supplementary Figure 11C**). When TUNEL assays were repeated with or without TNFα-challenge, apoptosis in response to such challenge was increased significantly only in NSNP (B1)- and stricturing (S, B2) CD-PDOs (**Supplementary Figure 11D**). All 3 subtypes of CD-PDOs showed increased apoptosis compared to TNFα-challenged healthy PDOs (**Supplementary Figure 11D**). Flow cytometry based TUNEL analyses were attempted, but not interpretable, because disruption of organoids was associated with artifacts.

Findings demonstrate that IDICD, but not S2FCD shows a higher rate of apoptosis and proliferation, and hence, higher cell turnover (see **Table 1**), which agrees with prior observations in the CD colon^62, 64–66^. Because cytokine-induced apoptosis appears to be functionally far more relevant than spontaneous apoptosis in causing barrier-dysfunction^67^ in IBD, results in TNFα-challenged conditions indicate that the CD-PDOs may have a wider range of defects in the milieu of inflammation.

### S2FCD and IDICD show subtype-defining phenomes that are therapeutically reversible

Next we asked if the predicted unique disease-defining phenotypes of CD-PDOs, i.e., immunodeficiency in the setting of infection in IDICD and senescence and genotoxic stress in S2FCD, are detected in the PDOs (**Figure 5A**). If so, such phenotypes could be tracked, and their reversal could be confirmed by rationally paired therapeutics.

**Figure 5.**
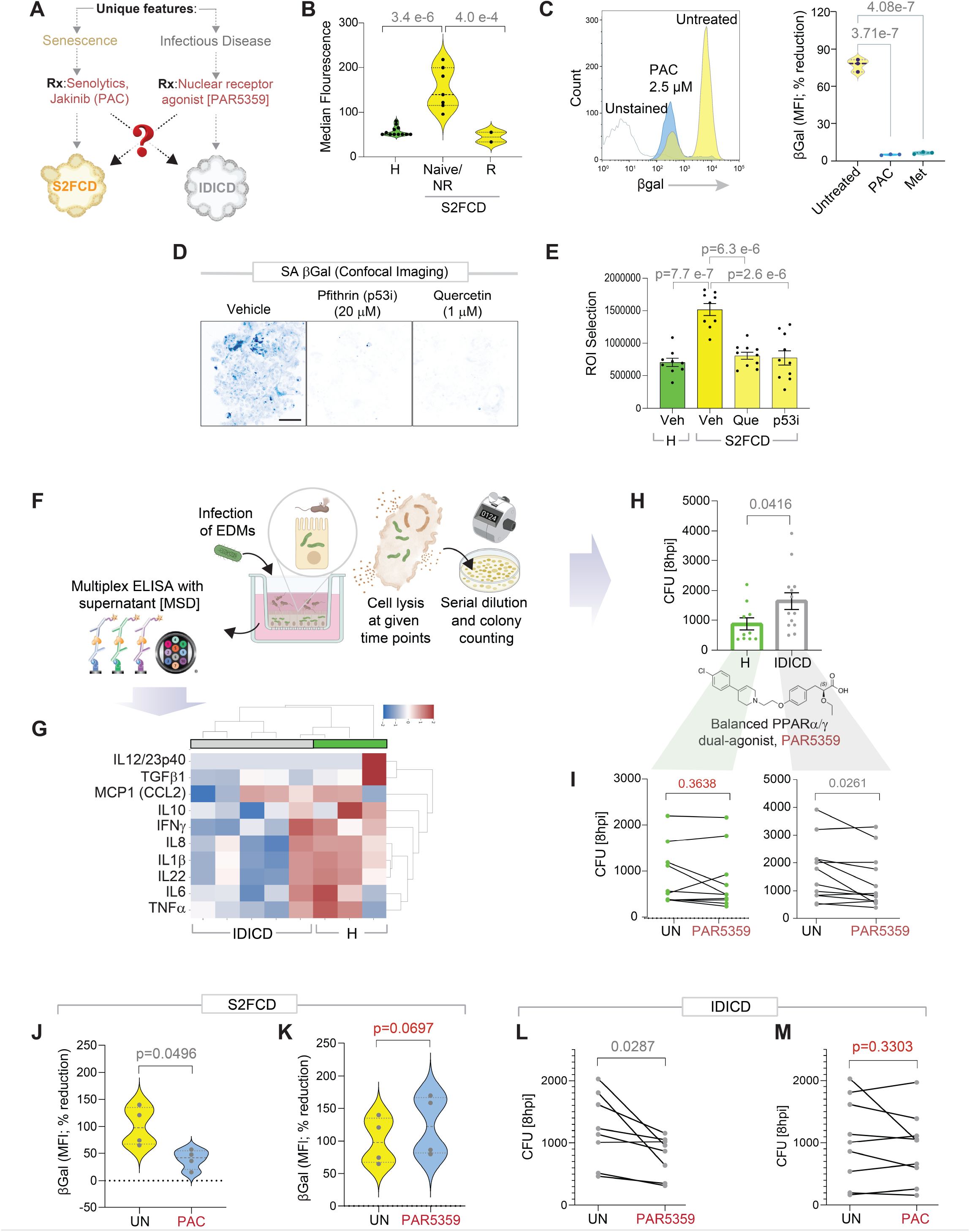
Genotyped-phenotyped CD-PDOs can serve as platforms for personalized therapeutics. **A.** Schematic outlines the approach towards validation and reversal through rationalized therapeutic pairing of the unique disease driver phenotypes predicted in each molecular subtype of CD (B-I). Subtype-paired therapeutics are also assessed for cross-over efficacy across the two molecular subtypes of CD (J-M). **B.** Violin plots show the median fluorescence intensity observed in healthy and CD-PDOs by flow cytometry [B- yellow cluster CD-PDOs separated into groups that responded (R) or not (NR) to anti-TNFα biologics or were naïve to that treatment]. Data represents 2-4 technical repeats on 4 healthy and 6 CD-PDOs. Statistical significance was assessed by one way ANOVA. **C.** Histogram (left) and violin plots (right) show the % changes in the median fluorescence intensity when CD- PDOs were treated with senotherapeutics (2.5 µM PAC, Pacritinib; 1 mM Met, Metformin). Statistical significance was assessed by one way ANOVA. **D-E**. Cellular senescence in PDOs was assessed using SPiDER βGal assays and subsequent analysis by immunofluorescence. Inverted images displayed (D) are representative of ∼10 fields/sample of max-projected z- stacks of CD-PDOs stained with *SPiDER*-SA-β-Gal. Bar graphs (E) display the quantification of staining. Scale bar = 100 µm. Statistical significance was assessed by one way ANOVA. **F.** Schematic outlines the 3 major steps in bacterial clearance assays, and the concomitant assessment of supernatants for cytokines by multiplexed ELISA. **G.** Heatmap displays the results of hierarchical agglomerative clustering of *AIEC*-LF82-challenged healthy and CD EDMs using the cytokine profiles determined by mesoscale (MSD). See **Supplemental Information 7**. **H.** Bar plots show the abundance of bacteria retained within healthy (H) and IDICD EDMs at 8 h after infection. Statistical significance was assessed by t-test. **I.** Line plots show the pre-(UN) and post-treatment (Rx) effect of a balanced PPARα/g dual agonist (1 µM PAR5359) on the abundance of bacteria at the 8h time point in healthy (left) and IDICD (right) EDMs. Statistical significance was assessed by t-test. **J-K**. Violin plots show the % change in median fluorescence intensity with PAC (J) or PAR5359 (K), as determined on S2FCD PDOs with the highest senescence in B. Statistical significance was assessed by t-test. **L-M**. Line plots show the untreated and effect of PAR5359 (L) or PAC (M) treatment on the abundance of bacteria at the 8h time point in IDICD EDMs with the highest bacterial load in H. Statistical significance was assessed by t-test. See **Supplemental Information 2** for subjects analyzed in each assay.

Compared to healthy PDOs, the S2FCD PDOs displayed increased cellular senescence, as determined using the established methods SA-βGal assay^68^; this phenotype was observed primarily in those who were biologic-naïve or those who were non-responder to anti-TNFα therapy (**Figure 5B**). Senescence was reversed using the senotherapeutic Pacritinib (PAC; **Figure 5C**), which inhibits JAK1/2 kinases that are prominent drivers of cellular senescence and its profibrogenic senescence-associated secretory phenotype (SASP)^69^. Metformin, which reverses numerous hallmarks of senescence^10^, was used as a positive control in these assays (Met; **Figure 5C**). Reduced senescence was also observed with the senolytic Quercetin-3-D-galactose (Que) and the senomorphic Pifithrin-α-p53 Inhibitor (p53i) in the CD-PDOs (**Figure 5D-E**).

In the case of IDICD, immune deficiency in the setting of infection was predicted the unique disease-defining phenotype (**Figure 2D**-*top left*). We infected EDMs prepared using healthy and IDICD-PDOs and analyzed them for their ability to mount a cytokine response and clear the infection (i.e., bacteria uptake and clearance assay; **Figure 5F-H**). For mimicking infection, we chose the pathogenic adherent invasive *Escherichia coli* strain-*LF82* (AIEC*-LF82*), which was isolated from CD patients^70^ and applied it to the apical surfaces of polarized EDMs using well-established protocols^22, 46^. Cytokine response was assessed by carrying out ultrasensitive, ELISA-based HTP proteome studies on the supernatants collected from infected EDMs. Compared to health EDMs, IDICD-EDMs showed impaired production of cytokines (**Figure 5G**). This defective cytokine response was only observed in the setting of infection (not at basal state without bacterial challenge) and only restricted to the IDICD, but not S2FCD-PDOs (**Supplemental Information 7**). Poor cytokine response in IDICD-EDMs was also accompanied by impaired clearance (i.e., more live bacteria recovered at 8 h post-infection; **Figure 5H**), indicating that CD-EDMs fail to clear bacteria and/or permit more replication. These findings agree with the prior findings of downregulated transcripts of multiple cytokines (**Figure 2D**-*bottom left*) and confirm the presence of an immune suppressed state in the setting of infection. We asked if impaired bacterial clearance in CD-EDMs can be reversed by a balanced dual agonist of PPARα/γ, PAR5359^71^, which has recently been shown to accelerate bacterial clearance in CD-patient derived PBMCs and ameliorate colitis in both DSS-induced chemical and *Citrobacter*-induced infectious colitis models^72^. We found that treatment of the CD-EDMs with PAR5359 significantly reduced the bacterial burden (**Figure 5I***-right*), with virtually no effect on healthy-EDMs (**Figure 5I**-*left*).

Although the phenotype of epithelial barrier dysfunction was not a prominent defect, and was observed primarily in the NSNP (B1) CD-PDOs, we asked if this too can be reversed. We prioritized two agents: The first is the probiotic drug *E coli* Nissle 1917 (*EcN;* **Supplementary Figure 12A**), which is safe and effective in maintaining remission equivalent to the gold standard mesalazine in patients with UC, but with dubious results in CD^73, 74^. The second is the post-biotic Hylak® Forte, which contains metabolic products (e.g., short-chain fatty acids [SCFA], amino acids, and vitamins) derived from commensal microbes (**Supplementary Figure 12B**). Both agents significantly increased the TEER in NSNP (B1) CD-EDMs, and in stricturing (B2) CD-EDMs.

### Subtype-personalized therapeutics lack cross-over efficacy

The surprising degree of internal consistency between the transcriptome, genome, and phenome, led us to ask if the molecular subtypes represent an opportunity for personalized therapeutics. If so, the phenotype-based pairing of therapeutics in one molecular subtype should lack efficacy when crossed over to the other subtype (**Figure 5A**). We compared head-to-head the efficacy of the lead drug-like candidates—PAC and PAR5359— on the most diseased CD-PDOs, i.e., those that were found to be most senescent (in **Figure 5B**) or most impaired in their ability to clear microbes (in **Figure 5H**). While PAC was effective in reversing senescence in S2FCD PDOs (**Figure 5J**), PAR5359 was not (**Figure 5K**). Similarly, while PAR5359 was effective in improving microbial clearance in IDICD PDOs (**Figure 5L**), PAC was not (**Figure 5M**). Findings suggest that subtype-specific therapeutic pairing may improve precision in targeting the disease-driving features within each subtype.

## CONCLUSION

The major discovery we report here is the identification of two distinct molecular subtypes of CD– IDICD and S2FCD-- based exclusively on the properties of the epithelial stem cells in the colon. The transcriptome, phenome, and even the underlying genome of each subtype is internally consistent and converge on a set of unique disease-driving attributes for each subtype. Proof-of-concept studies demonstrate that molecular subtyping of CD-PDOs, as opposed to the current clinical classification system, can enable personalized therapeutics to precisely reverse the unique disease-driving attribute of that subtype with little or no cross-over efficacy. These findings reveal the potential for genotyped-phenotyped CD-PDOs as tools for implementing personalized medicine. They also represent a paradigm shift in how we classify CD from clinical patterns to dysregulated molecular pathways that directly suggest therapeutic interventions.

There are three major impacts of these findings.

### A CD molecular classification that reconciles genome, transcriptome, and phenome

Finding a robust internal consistency between genome, transcriptome, proteome (cytokines) and phenome that overcame the heterogeneity of clinical presentation and cohort composition is striking. The IDICD subtype shows impaired pathogen clearance and insufficient cytokine response. This subtype frequently harbors the major CD risk-alleles of NOD2 and ATG16L1 and shows skewed differentiation in the PDOs. The S2FCD subtype, is characterized by cellular senescence, genotoxic stress, and displays a profibrotic transcriptome. The genome of this CD subtype shows higher mutations in the YAP1-IL-18 pathway, which has recently been implicated in telomere instability-associated tissue inflammation that originates in the CD colonic epithelium^38^. These findings are consistent with prior work showing that DNA damage serves as a trigger for senescence and inflammation in CD^75^. Up until now, the contributions of genetics as a determinant in the various clinical behaviors of CD have remained unclear^3^; the alignment of genetics with the transcriptome and phenome within each molecular subtype suggests that the molecular subtypes (revealed here) could be a more meaningful way to classify patients because such classification is linked to driver phenotypes of the disease with therapeutic implications. Finally, that perianal disease mimics penetrating (B3) CD at a fundamental molecular level is another important insight, which should be exploited in determining management strategies for these patients.

### Molecular classification that informs the choice of therapeutics

Efforts at the successful development of CD therapeutics have stumbled because of our lack of understanding of the factors that affect disease heterogeneity and evolution. That a vast majority of the patients belong to the IDICD molecular subtype may explain the observed futility^16^ of the currently FDA-approved immunosuppressive drugs as agents to maintain remission in B3-penetrating clinical subtype (which, belong to the IDICD molecular subtype). Our proof-of-concept studies also reveal that CD-PDOs could be used to directly test drug efficacy in a personalized treatment approach; PDOs could be classified into one of the two major molecular subtypes and subsequently tested for therapeutic efficacy using novel or clinically approved drugs within weeks after derivation. In doing so, CD-PDOs may fill the gap between imperfect cell/animal models, limitations of CD GWAS and clinical trials and allow personalized therapy selection. For example, the pan-JAK-inhibitor Tofacitinib has succeeded in Phase III trials for UC, with conflicting results in CD^76^, and our findings suggest that targeted trials on the S2FCD molecular subtype using JAK1/2-selective inhibitors (e.g., Upadacitinib, Filgotinib or Baricitinib) may have demonstrable efficacy. As for IDICD, the strategy of improving microbial clearance using the balanced dual-PPARα/γ agonism with PAR5359 was identified using an AI-guided network transcriptomics approach^20, 72^, which is predicted to protect the gut mucosal barrier in IBD and represents a new class of therapy in IBD yet to enter clinical trials.

### A possible framework for benchmarking PDOs as models to study altered epithelial biology in the inflamed gut

It is important to note that PDOs arise from regenerating adult stem cells, and that ultimately, the disease-driving transcriptome and phenome in the PDOs reflect the consequences of genetic, epigenetic, and feedback regulatory events also from the cell signaling circuits. Recapitulating the active diseased epithelium, despite being removed for prolonged duration (∼weeks to months) from the dysbiotic lumen and the inflamed *in vivo* conditions suggests that these adult stem cell-derived PDOs may retain the ‘memory’ of the disease state. Objective benchmarking metrics (i.e., gene expression in PDOs vs CD epithelium; **Figure 2C**; **Supplementary Figure S1**) revealed that retention of such ‘memory’ may require growth conditions that have proven track record of reproducibility between laboratories^23^, and may require the use of colon-crypt derived adult stem cells (instead of reprogrammed iPS cells). This disadvantage of iPS-derived organoids as models for IBD was predicted intuitively^77^, but remained unproven until now.

## LIMITATIONS OF THE STUDY

Although bulk RNA seq is insightful, and allowed us to profile a larger cohort, future single-cell studies are required to obtain a higher resolution on cell-type specific aberrant processes in each molecular subtype of CD. A side-by-side comparison of PDO vs. its tissue of origin was not attempted; prior studies have shown cell type composition and > 90% similarities in protein-coding genes between CD tissue and CD-PDOs^8^. Despite these shortfalls, it is noteworthy that our findings in CD-PDOs by bulk seq mirror several key claims of another article in pre-print that used single cell sequencing on primary epithelial cells from CD colons^10^. Because the PDOs retained transcriptome and phenome during prolonged cultures, the epigenetic events that preserve the memory of the *in vivo* diseased state is an interesting avenue that was not explored here. The current study also did not assess the ileum or the microbiome; such studies from the same patient are expected to give a more complete picture. For example, we expect to see more dysbiosis in the IDICD subtype compared to the S2FCD subtype.

## ACKNOWLEDGEMENTS

This work was supported by the Leona M. and Harry B. Helmsley Charitable Trust (to PG and SD). Other sources of support include-- National Institutes for Health (NIH) grants AI141630 (to P.G), DK107585, R56 AG069689 and DiaComp Pilot and Feasibility award (to SD), R01-GM138385, Padres Pedal the Cause/C3 Collaborative Translational Cancer Research Award (San Diego NCI Cancer Centers Council [C3] #PTC2017) (to DS). PG, SD, and DS were also supported by UG3TR003355, UG3TR002968 and R01-AI155696. B.S.B. was supported by NIH K23DK123406 and P30DK120515. GDK was supported through The American Association of Immunologists Intersect Fellowship Program for Computational Scientists and Immunologists. SRI was supported by the postdoctoral fellowship grant from NIH (3R01DK107585-02S1). J. E. was supported by an NCI/NIH-funded Cancer Biology, Informatics & Omics (CBIO) Training Program (T32 CA067754) and a Postdoctoral Fellowship from the American Cancer Society (PF-18-101-01-CSM). The content is solely the responsibility of the authors and does not necessarily represent the official views of the Helmsley Charitable Trust or the National Institutes of Health. This publication includes data generated at the UC San Diego IGM Genomics Center utilizing an Illumina NovaSeq 6000 that was purchased with funding from a National Institutes of Health SIG grant (#S10 OD026929). We are grateful to Helen Le and Jennifer Neill (UCSD IBD Center), Donald Pizzo (UCSD Pathology Histologic Biomarkers Core), Ying Jones (UC San Diego Electron Microscopy Core Facility), Joshua Alcantara, Vanae Goheen-Holland, Kevin Vega, Uddeep Chaudhary and Amanraj Claire (UC San Diego HUMANOID™ Center of Research Excellence) for technical and logistical support. We also thank student intern Poorvi Saini and Interval Bio for assistance with the processing of genotyping datasets.

## AUTHOR CONTRIBUTIONS

S.D. and P.G. conceptualized, supervised, administered the project and acquired funding to support it. C.T., A.G.F, G.D. K., I.I.S., S.R.C., R.P., M.F., P.M., D.L.S., S.D., P.G. were involved in organoid isolation, culture and their use in various experiments, data curation and formal analysis. Computational analyses were carried out by S.T under the supervision of P.G and D.S. D.S. provided all computational software. H.L.N., W.J.S. and B.S.B provided key resources for human subjects and were responsible for the selection and enrolling of patients into this study. C.T, A.G.F, S.D., and P.G. prepared figures for data visualization, wrote the original draft. C.T, A.G.F, B.S.B., D.S., S.D., P.G., I.M.S., and G.D.K. reviewed and edited the draft. All co-authors approved the final version of the manuscript.

## SUPPLEMENTAL INFORMATION

**Supplemental Information 1:** Excel datasheet with the list of healthy and CD patients who were enrolled into this study and their demographic information.

**Supplemental Information 2:** Excel datasheet with the list of healthy and CD PDOs that were analyzed in various functional assays.

**Supplemental Information 3:** Excel datasheet with the list of healthy and CD PDOs that were analyzed by RNA Seq.

**Supplemental Information 4:** Excel datasheet with the list of differentially regulated genes in CD-PDOs and their reactome pathway analyses.

**Supplemental Information 5:** Excel datasheet with gene list and rsIDs used for genotype analysis on CD- PDOs.

**Supplemental Information 6:** Excel datasheet with QPCR primer sequences

**Supplemental Information 7:** Excel datasheet with source data for cytokine analysis by MSD.

## MATERIALS AND METHODS

### KEY RESOURCES TABLE

| Reagent or Resource | Source | Identifier |
| --- | --- | --- |
| **Antibodies** |  |  |
| Recombinant Anti-Collagen I Antibody | Abcam | Cat# ab270993 |
| Anti-β-catenin Antibody (E-5) | Santa Cruz | sc-7963 |
| E-cadherin Antibody (H-108) | Santa Cruz | sc-7870 |
| Recombinant Anti-Ki67 antibody [SP6] | Abcam | ab16667 |
| P-Histone H2A.X (Ser 139) | Santa Cruz | sc-517348 |
| Lysozyme Polyclonal Antibody | Invitrogen | PA5-16668 |
| MUC2 Monoclonal Antibody (996/1) | Invitrogen | MA5-12345 |
| Occludin Monoclonal Antibody (OC-3F10) | Invitrogen | 33-1500 |
| Vimentin Monoclonal Antibody (3H9D1) | Proteintech | 60330-1-Ig |
| ZO-1 antibody [N2C1], Internal | GeneTex | GTX108627 |
| Goat Anti-Rabbit Alexa Fluor 488 Antibody | Abcam | Cat# 150077<br>RRID: AB_2630356 |
| Goat Anti-Mouse Alexa Fluor 488 Antibody | Invitrogen | A11001 |
| Goat Anti-Rabbit Alexa Fluor 594 Antibody | Invitrogen | A11012 |
| Goat Anti-Rabbit Alexa Fluor 488 Antibody | Abcam | Cat# 150077<br>RRID: AB_2630356 |
| **Bacterial strains** |  |  |
| Adherent-Invasive *E. coli* LF82 | Arlette Darfeuille-Michaud | NA |
| **Chemicals, peptides, and recombinant proteins** |  |  |
| MEM | Corning | Cat# MT 10-010-CV |
| Collagenase Type I | Thermo Fisher | Cat# 17100017 |
| Collagenase Type II | Thermo Fisher | Cat# 17101015 |
| Collagenase Type IV | Thermo Fisher | Cat# 17104019 |
| HBSS with Ca<sup>2+</sup>/Mg<sup>2+</sup> | Thermo Fisher | Cat# 14-025-092 |
| HBSS without Ca<sup>2+</sup>/Mg<sup>2+</sup> | Thermo Fisher | Cat# 14-175-095 |
| MEM Non-Essential Amino Acids | Thermo Fisher | Cat# 11-140-050 |
| Sodium Pyruvate | Sigma-Aldrich | Cat# S8636 |
| Ciprofloxacin Hydrochloride | Corning | Cat# 61-277-RF |
| Antibiotic Antimycotic | Sigma-Aldrich | Cat# A5955 |
| Trypsin (2.5%) | Thermo Fisher | Cat# 15090046 |
| Zinc Formalin | Fisher Scientific | Cat# 23-313096 |
| Xylene | VWR | Cat# XX0060-4 |
| Hematoxylin | Sigma-Aldrich Inc | Cat# MHS1 |
| Ethanol | Koptec | Cat# UN1170 |
| Sodium Citrate | Sigma-Aldrich | Cat# W302600 |
| DAB (10x) | Thermo Fisher | Cat# 1855920 |
| Stable Peroxidase substrate buffer (10x) | Thermo Fisher | Cat# 34062 |
| 3% Hydrogen Peroxide | Target | Cat# 245-07-3628 |
| Horse Serum | Vector Labs | Cat# 30022 |
| Paraformaldehyde 16% Solution, EM Grade | Electron Microscopy Sciences | Cat# 15710 |
| 100% Methanol | Supelco | Cat# MX0485 |
| Glycine | Fisher Scientific | Cat# BP381-5 |
| Bovine Serum Albumin | Sigma-Aldrich | Cat# A9647-100G |
| Triton-X 100 | Sigma-Aldrich | Cat# X100-500ML |
| Prolong Glass | Invitrogen | Cat# P36984 |
| Nail Polish (Rapid Dry) | Electron Microscopy Sciences | Cat# 72180 |
| Gill Modified Hematoxylin (Solution II) | Millipore Sigma | Cat# 65066-85 |
| Histogel | Thermo Scientific | Cat# HG4000012 |
| TrypLE Select | Thermo Scientific | Cat# 12563-011 |
| Advanced DMEM/F-12 | Thermo Scientific | Cat# 12634-010 |
| HEPES Buffer | Life Technologies | Cat# 15630080 |
| Glutamax | Thermo Scientific | Cat# 35050-061 |
| Penicillin-Streptomycin | Thermo Scientific | Cat# 15140-122 |
| Matrigel | Corning | Cat# 354234 |
| DPBS | Thermo Scientific | Cat# 14190-144 |
| Ultrapure Water | Invitrogen | Cat# 10977-015 |
| EDTA | Thermo Scientific | Cat# AM9260G |
| Fetal Bovine Serum | Sigma-Aldrich | Cat# F2442-500ML |
| Cell Recovery Solution | Corning | Cat# 354253 |
| Sodium Azide | Fisher Scientific | Cat# S227I-100 |
| Cyto-Fast Fix/Perm Buffer Set | BioLegend | Cat# 426803 |
| FITC-Dextran | Sigma-Aldrich | Cat# FD10S |
| Ethyl alcohol, pure | Sigma-Aldrich | Cat# E7023 |
| TRI Reagent | Zymo Research | Cat# R2050-1-200 |
| Metformin | Sigma | Cat# D150959-5G |
| Pacritinib | Selleck Chem | Cat# S8057 |
| Quercetin | Sigma Aldrich | Cat# Q4951 |
| TP53 Inhibitor (p53i; Pifithrin-α-HBr) | Adipogen Life Science® | Cat# AG-CR1-0004-M005 |
| Hylak Forte™ | Amazon.com |  |
| **Critical commercial assays** |  |  |
| BrdU Cell Proliferation ELISA Kit (colorimetric) | Abcam | Cat# ab126556 |
| DNA/RNA oxidative damage ELISA kit | Cayman Chemical, USA | Cat # 589320 |
| SPiDER-βGal | Dojindo | Cat# SG02-10 |
| TUNEL Assay Kit - BrdU-Red | Abcam | Cat# ab66110 |
| Quick-RNA MicroPrep Kit | Zymo Research | Cat# R1051 |
| Quick-RNA MiniPrep Kit | Zymo Research | Cat# R1054 |
| HRP Horse Anti-Rabbit IgG Polymer Detection Kit | Vector Laboratories | Cat# MP-7401 |
| **Oligonucleotides** |  |  |
| 2x SYBR Green qPCR Master Mix | Bimake | Cat# B21203 |
| qScript cDNA SuperMix | Quanta Biosciences | Cat# 95048 |
| **Software and algorithms** |  |  |
| ImageJ | ImageJ | RRID:SCR_003070 |
| GraphPad Prism | GraphPad Prism | RRID:SCR_002798 |
| MSD® DISCOVERY WORKBENCH 4.0 | MSD |  |
| LAS AF Software | LAS AF Software |  |
| QuantStudio Design & Analysis Software | QuantStudio Design & Analysis Software |  |
| CIBERSORTx | CIBERSORTx |  |
| FlowJo | Flow Jo V10, BD BioSciences | RRID:SCR_008520 |
| **Other** |  |  |
| 6-well Tissue Culture Plate | Genesee Scientific | Cat# 25-105 |
| 12-well Tissue Culture Plate | CytoOne | Cat# CC7682-7512 |
| Transwell Inserts (6.5 mm, 0.4 um pore size) | Corning | Cat# 3470 |
| Cell Scraper | Millipore Sigma | Cat# C5981-100EA |
| Millicell EZ Slide 8-Well Chamber | Millipore Sigma | Cat# PEZGS0816 |
| Trypan Blue Stain | Invitrogen | Cat# T10282 |
| 70 um Cell Strainer | Thermo Fisher Scientific | Cat# 22-363-548 |
| Noyes Spring Scissors - Angled | Fine Science Tools | Cat# 15013-12 |
| gentleMACS™ C Tubes | Miltenyi Biotec | Cat# 130-093-237 |
| DAPI | Invitrogen | Cat# D1306<br>RRID: AB_2629482 |
| Phalloidin, Alexa Fluor 594 | Invitrogen | Cat# A12381<br>RRID: AB_2315633 |
| Propidium Iodide | Invitrogen | Cat# V-13245 B |
| Countess II Automated Cell Counter | Thermo Fisher Scientific | AMQAX1000 |
| Epithelial Volt-Ohm (TEER) Meter | Millipore | MERS00002 |
| Automated TEER measurement system [REMS AutoSampler] | World Precision Instruments (WPI) |  |
| MESO QuickPlex SQ 120 | MSD |  |
| Leica TCS SPE Confocal | Leica Microsystems | TCS SPE |
| Power Pressure Cooker XL | Tristar Products |  |
| Canon Rebel XS DSLR | Canon |  |
| MiniAmp Plus Thermal Cycler | Applied Biosystems | Cat# A37835 |
| QuantStudio5 | Applied Biosystems | Cat# A28140<br>RRID:SCR_020240 |
| Light Microscope (brightfield images) | Carl Zeiss LLC | Axio Observer,<br>Inverted; 491917-0001-000 |
| Fisherbrand™ 150 Handheld Homogenizer | Fisher Scientific | Cat# 15340168 |
| Spark 20M Multimode Microplate Reader | Tecan |  |
| NanoQuant Infinite M200 | Tecan |  |
| Guava® easyCyte Benchtop Flow Cytometer | Millipore | Guava easyCyte 6 2L |
| MESO QuickPlex instrument | Mesoscale Discovery Inc. | SQ 120 |
| gentleMACS™ Dissociator | Miltenyi Biotec | Cat# 130-093-235 |

### RESOURCE AVAILABILITY

#### Lead Contact

Further information and requests for resources and reagents should be directed to and will be fulfilled by the lead contact, Pradipta Ghosh.

#### Materials availability

This study has generated organoid biobank, RNA and DNA from the organoids. These materials can only be accessed through proper material transfer and patented technology (Das & Ghosh) agreement following the guidelines of the University of California, San Diego.

#### Data and code availability

Source data are provided with this paper. All data is available in the main text or the supplementary materials. All source data supporting the graphs in this manuscript will be uploaded and made publicly available upon publication. All RNA sequence data and processed data in this manuscript have been deposited at Gene Expression Omnibus NCBI GEO (GSE192819). Publicly available datasets used: GSE16879; GSE115390; E- MTAB-7604.

### EXPERIMENTAL MODEL AND SUBJECT DETAILS

#### Human Subjects

For generating healthy and CD patient-derived organoids (PDOs), patients were enrolled for colonoscopy as part of routine care for the management of their disease from the University of California, San Diego IBD-Center, following a research protocol compliant with the Human Research Protection Program (HRPP) and approved by the Institutional Review Board (Project ID# 1132632: PI Boland and Sandborn). Histologically normal healthy colon samples were collected from patients presenting for screening colonoscopy or undergoing the procedure for making the diagnosis of irritable bowel syndrome. Each participant provided a signed informed consent to allow for the collection of colonic tissue biopsies for research purposes to generate 3D organoids. Isolation and biobanking of organoids from these colonic biopsies were carried out using an approved IRB (Project ID # 190105: PI Ghosh and Das) that covers human subject research at the UC San Diego HUMANOID Center of Research Excellence (CoRE). For all the deidentified human subjects, information including age, gender, and previous history of the disease, was collected from the chart following the rules of HIPAA. The study design and the use of human study participants was conducted in accordance to the criteria set by the Declaration of Helsinki.

#### Isolation of enteroids from colonic specimens of healthy and Crohn’s Disease subjects

Intestinal crypts, comprised of crypt-base columnar (CBC) cells, were isolated from human colonic tissue specimens using the previously published paper^20–22, 47^. In brief, intestinal crypts were dissociated from tissues by digesting with collagenase type I (2 mg/mL solution containing gentamicin 50 µg/mL). The plate was incubated in a CO_2_ incubator at 37°C, mixing every 10 min with vigorous pipetting in-between incubations, while monitoring the release of single epithelial units from tissue structures by light microscopy. To inactivate collagenase, wash media (DMEM/F12 with HEPES, 10% FBS) was added to cells, filtered through a 70 μm cell strainer, centrifuged at 200 g for 5 min and then the supernatant was aspirated, leaving behind a cell pellet. The number of viable intestinal stem cells was determined by the Trypan Blue Exclusion method using Countess II Automated Cell Counter. Epithelial units were resuspended in Matrigel and 25 μl of cell-matrigel suspension was added to the wells of a 12-well plate on ice and incubated upside-down in a 37°C CO_2_ incubator for 10 min, which allowed for polymerization of the Matrigel. After 10 min of incubation, 1000 μL of 50% conditioned media, prepared from L-WRN cells with Wnt3a, R-Spondin and Noggin, ATCC® CRL-3276™^19^ with a GI-organoid media cocktail 1 (purchased from the UC San Diego HUMANOID™ Center of Research Excellence), 10 μM Y27632 and 10 μM SB431542. The medium was changed every 2 days and the enteroids were either expanded or frozen in liquid nitrogen for biobanking. The number of PDOs varied between patient samples, with some tissues rendering hundreds-to-thousands of organoids, whereas others yielded only tens-to-hundreds primary organoids. Organoids from patients with active inflammation took longer to establish than those without inflammation.

#### Preparation of Enteroid-derived monolayers (EDMs)

EDMs were prepared by dissociating single cells from enteroids and plated either in 24-well or 96-well transwell with a 0.4 μm pore polyester membrane coated with diluted Matrigel (1:40) in 5% conditioned media as done before^20–22, 47^. The single-cell suspension was seeded at a density of approximately 2×10^5^ cells/well (in case of 24-well) or 8×10^4^ cells/well (in case of 96-well) and EDMs were differentiated for 2-3 days in 5% conditioned media. The media was changed every 24 h and monitored under a light microscope to evaluate the EDM generation and quality. As expected, the expression of EDMs showed a significant reduction of the stemness marker *LGR5* in EDMs^21, 25, 47^.

### METHOD DETAILS

#### Experimental methods

##### Quantitative assessment of organoid morphology by Imaris

.LIF files were first converted into native IMARIS format (.ims). Then a spots filter and surface filter were created. This filter is used as a batch function on all processed images. Finally, a cell object is created where broken fragments of single organoids are stitched together manually. Upon manually completion specific measurements are exported from IMARIS to GraphPad Prism for further analysis and for visualization as graphs.

##### TUNEL assay

The baseline level and the effect of TNFα on apoptosis between healthy and CD organoids was quantified using the TUNEL Assay Kit via immunofluorescence (see Key Resource Table). Organoids were seeded at a density of 5000 cells/well on a layer of Matrigel in an 8-well chamber slide and cultured for 4 days in growth media. TNFα (100 ng/mL) treatment was applied for 24 h prior to fixation (day 3). Organoids were fixed in 4% paraformaldehyde (PFA) at room temperature for 30 min and quenched with 30 mM glycine for 5 min. Subsequently, samples were processed as per manufacturer’s specification, i.e., they were permeabilized and blocked for 1 h using blocking buffer (2 mg/mL BSA and 0.1% Triton X-100 in PBS), washed with PBS and then treated with 100 µL of DNA labelling solution (diluted 1:1) in ultrapure water, and incubated subsequently in a dark humidified incubator for 1 h at 37°C. A negative control DNA labelling solution, which substituted the TdT labelling enzyme for ultrapure water, was used to determine assay background and noise. Samples were washed in PBS, treated with 100 µL of antibody solution (constituted according to the manufacturer’s specifications) supplemented with DAPI (1:500 dilution) and then incubated in the dark for 30 min at room temperature. Samples were washed in ultrapure water, mounted using Prolong Glass prior to application of coverslips (no. 1 thickness) and sealed. Slides were imaged immediately by confocal microscopy within 6 h of staining as per manufacturers recommendation.

##### Quantification of TUNEL assays

Z-stack images were acquired by successive 3 µm depth Z-slices of organoids using confocal microscopy. Maximum intensity projection of the stack of images were analyzed using the FIJI (ImageJ) Plugin, ‘RGB Measure’ to quantify the integrated density of the BrdU signal (red/594 nm channel). Total area of enteroids in a field was determined by converting DAPI (blue) channel to a binary image and thresholded to determine nuclear boundaries. The level of apoptosis was quantified by comparing apoptotic-positive cell signal (Brdu-Red) relative to the area of DAPI-positive cell signal. A blank subtraction, using the negative control, was done to normalize values across independent experiments. The effect of TNFα on each patient-derived organoid line was determined by fold change relative to its untreated sample counterpart. All images were processed on ImageJ software (NIH) and assembled into figure panels using Photoshop and Illustrator (Adobe Creative Cloud).

##### Assessment of cell proliferation by an ELISA-based BrdU incorporation assay

The level of proliferation was quantified with a BrdU Cell Proliferation ELISA kit (see Key Resource Table). Organoids were seeded at a density of 3500 cells/well on a layer of Matrigel in 96-Well plate and cultured for 3 days in growth media. BrdU (1x) was added and allowed to incorporate into the organoids for 24 h prior to fixation on day 4. Cells were fixed according to the manufacturer’s instructions. Optical Density (OD) was measured using a 450 nm spectrophotometer microplate reader. Each patient cell line had negative BrdU control wells for the determination of background signal. The average negative BrdU control OD was subtracted from a patients average BrdU-incorporated OD to obtain a normalized OD value. Fold change was determined relative to a single healthy PDO, kept constant through all assays, for all samples including other healthy PDOs.

##### Assessment of cell proliferation by Ki67 particle analysis

Z-stack images were acquired by successive 3 µm depth Z-slices of Ki67 stained organoids using confocal microscopy. From the acquired maximum intensity projection images, the DAPI channel was converted to a grayscale image. A gaussian blur filter of σ (Radius): 1 was applied to reduce nuclear noise. Manual thresholding was applied to convert grayscale image into a binary: black and white image; using the FIJI Default settings and Over/Under against a dark background. Overlapping particles were separated using the watershed method and particles were analyzed with the requirements; size: 5-infinity µ^2, circularity: 0.4-1.0 and edge particles excluded. The auto generated DAPI particle outline was used as a boundary to manually count particles that contained positive Ki67 signal. The #Ki67 positive cells/total # of cells was used to generate the % of Ki67 positive cells. Approximately 10 random fields, that were representative of the overall staining observed in the samples were imaged per PDO.

##### Measurement of oxidative DNA/RNA damage

The amount of oxidative DNA damage in healthy and CD EDMs was quantified using a commercial kit (see Key Resource Table) according to the manufacturer’s instructions and previously published papers by others^78–80^ and us^81, 82^. Briefly, supernatant from EDMs was used to detect oxidized guanine species: 8-hydroxy-2′- deoxaguanosine from DNA, and 8-hydroxyguanine from either DNA or RNA.

##### Estimation of Paneth:Goblet cell ratio by confocal imaging of cell markers

Fluorescent Z-stack images of lysozyme (a *bona-fide* marker of Paneth cells) and muc2 (a *bona-fide* marker of goblet cell) stained organoids were acquired by successive 1 μm depth Z-slices of EDMs in the desired confocal channels of Leica TCS SP5 Confocal Microscope as done previously^47^. Fields of view that were representative of a given transwell were determined by randomly imaging 3 different fields. Z-slices of a Z-stack were overlaid to create maximum intensity projection images; all images were processed using FIJI (Image J) software. All images were processed on ImageJ software (NIH) and assembled into figure panels using Photoshop and Illustrator (Adobe Creative Cloud).

##### Measurement of LYZ: MUC2 ratio

From the acquired maximum intensity projection images both lysozyme (Red/594 nm channel) and MUC2 (green/488 nm channel) were converted to grayscale images. Particles were quantified, with the following thresholds-- size: 0-infinity µ^2^, circularity: 0.0-1.0 and edge particles were included to independently determine the total area positive for lysozyme and MUC2 signals. DAPI particle analysis was carried out the same as done for Ki67 particle analysis. Total area of cell-specific marker/total # of cells was used to generate a normalized value of area per cell for each marker, lysozyme and MUC2. Approximately 5-10 random fields that were representative of the staining were imaged per PDO sample. All images were processed on ImageJ software (NIH) and assembled into figure panels using Photoshop and Illustrator (Adobe Creative Cloud).

##### Embedding of organoids in HistoGel™

Healthy and CD colonic organoids were embedded in histogel as done previously^83^. Briefly, mature organoids after 7-dasys of culture in 6-Well plates were fixed in 4% PFA at room temperature for 30 min and quenched with 30 mM glycine for 5 min. After washing with PBS, organoids were resuspended in PBS and stained using Gill’s hematoxylin for 5 min for ease during embedding in paraffin blocks and visualization during and after sectioning. Excess hematoxylin was removed, and organoids were resuspended in HistoGel™ and centrifuged at 65°C for 5 min. HistoGel™ embedded organoid pellets were cooled to room temperature and stored in 70% ethanol at 4°C until ready for embedding in paraffin blocks. FFP-embedded organoid sections were cut at a setting of 4 µm thickness and fixed on to microscope slides for H&E staining.

##### Immunofluorescence of FFPE organoids

Sections of FFP-embedded healthy- and CD- PDOs were deparaffinized, rehydrated and underwent antigen retrieval immersed in Sodium Citrate buffer (pH 6.0) and boiled at 100°C inside a pressure cooker for 3 min. Once sections returned to room temperature, samples were washed in DI water and then permeabilized and blocked for 2 h using an in-house blocking buffer (2 mg/mL BSA and 0.1% Triton X-100 in PBS), as described previously^83, 84^. Primary antibodies [see Key Resource Table] were diluted in blocking buffer and incubated overnight at 4°C. Secondary antibodies were diluted in blocking buffer and allowed to incubate for 2 h in the dark. Antibody dilutions are listed in the Supplementary Key Resource Table. ProLong Glass was used as a mounting medium. Coverslips (No.1 thickness) were applied to slides to seal and stored at 4°C until imaged.

##### Ultrastructural analyses of patient-derived organoids by electron microscopy

Organoid pellets were fixed with 2% Glutaraldehyde in 0.10 M cacodylate buffer and further postfixed in 1% OsO4 in 0.10 M cacodylate buffer for 1 h on ice. Organoids were stained with 2% uranyl acetate for 1 hr on ice, following which they were dehydrated in a graded series of ethanol (50-100%) while remaining on ice. Organoids were then subjected to 1 wash with 100% ethanol and 2 washes with acetone (10 min each) and embedded with Durcupan. Sections were cut at 60 nm on a Leica UCT ultramicrotome and picked up on 300 mesh copper grids; different planes were obtained from consecutive 10 µm cuts. Sections were post-stained with 2% uranyl acetate for 5 min and Sato’s lead stain for 1 min. Images were acquired using a JEOL 1400 plus microscope equipped with a bottom-mount Gatan OneView (4k x 4k) camera.

##### Immunofluorescence imaging of epithelial tight junctions in EDMs

The media from apical and basolateral compartments of all EDMs were removed, washed 3 times with room temperature PBS, fixed with ice-cold 100% methanol at -20°C for 20 min. Afterward, methanol was removed and washed with blocking buffer (0.1% Triton TX-100, 2 mg/mL BSA diluted in PBS) to permeabilize EDMs and to incubate with the following primary antibodies overnight: ZO-1 (1:500) and Occludin (1:500). Primary antibodies were removed and washed with PBS 3 times for 5 min each time; after which the following secondary antibodies were added for 2 h: Alexa Fluor 594 conjugated goat anti-rabbit IgG, Alexa Fluor 488 conjugated goat anti-mouse IgG and DAPI. Secondary antibodies were removed and washed with PBS 3 times for 5 min each time. To preserve fluorescence, monolayers were treated with Prolong Gold antifade reagent and stored at 4°C until imaged. A confocal Microscope (Leica SPE) with a 40x objective lens was used to image the stained EDMs.

##### Measurement of transepithelial electrical resistance (TEER)

Two different methods in low- (LTP) and high-throughput (HTP) modes were used for the measurement of TEER as described before with some modifications^22^. LTP assessment of TEER was carried out manually in 24-well transwell plates. TEER was measured at 24 h, and 48 h, following monolayer seeding using STX2 electrodes with digital readout by EVOM2 (WPI). HTP assessment of TEER was carried out in an automated manner in 96- well transwell plates. TEER was measured using the REMS AutoSampler (WPI) automated TEER measurement system (see Key Resource Table). A WPI REMS-96C recording electrode was used to record TEER, which is compatible for use with a 96-well plate from Corning. The REMS-96C recording electrode was sterilized in 70% ethanol, followed by a rinse in PBS, and subsequently in media. The REMS-96C apical electrode was calibrated to measure TEER approximately 1 mm above the transwell membrane. Transwell-read time set to 5 sec/well. Once set-up is complete, the plate is removed from incubator and TEER is measured directly afterwards. To mitigate TEER artifacts due to temperature fluctuations, the same read sequence is repeated every subsequent read. TEER recorded by REMS AutoSampler were saved as .txt files; raw TEER values (in Ωs), are converted to normalized TEER values by Raw TEER in ohms (Ω) x surface area of transwell in cm2 = ohms. cm2 (SA=0.143 cm2 for 96-well and 0.33 cm2 for 24-well).

##### Assessment of barrier permeability of EDMs using FITC-dextran

EDMs were grown for 48 h in 5% CM on 96-well transwells and TEER was monitored by a pair of REMS-96C recording electrodes using the WPI automated TEER Measurement System. After 48 h of growth, FITC-dextran (10 kD) was added to the apical side at a 1:50 dilution in 5% conditioned media. After 1 h of incubation with FITC-dextran, 50 µl of the basolateral supernatant was transferred to an opaque-black 96-well plate. Fluorescence was measured using excitation/emission 485 nm/535 nm with a Spark 20M Multimode Microplate Reader (see Key Resource Table).

##### RNA isolation

Organoids and monolayers were lysed using 200 ul of RNA lysis buffer followed by RNA extraction per Zymo Research Quick-RNA MicroPrep Kit instructions (see Key Resource Table). Tissue samples were lysed in TRI-Reagent and RNA was extracted using Zymo Research Direct-zol RNA Miniprep.

##### Quantitative (q)RT-PCR

Organoid and monolayer gene expression was measured by qRT-PCR using 2x SYBR Green qPCR Master Mix. cDNA was amplified with gene-specific primer/probe set and qScript cDNA SuperMix (5x). qRT-PCR was performed with the Applied Biosystems QuantStudio 5 Real-Time PCR System. Cycling parameters were as follows: 95 °C for 20 s, followed by 40 cycles of 1 s at 95 °C and 20 s at 60 °C. Primers used in qRT-qPCR were previously validated in similar studies^85–87^ and primer sequences are stated in the **Supplemental Information 6.** All samples were assayed in triplicate and eukaryotic 18S ribosomal RNA was used as a reference.

##### DNA isolation

Organoid pellets were lysed using Genomic Lysis Buffer followed by DNA extraction per Zymo Research Quick-DNA Microprep Kit (see Key Resource Table).

##### Genomic analysis of patient-derived organoids

A list of ∼154K SNPs associated with IBD was created by searching published papers^88–90^, Clinvar and IBD genetics databases. A custom primer panel was developed by Tecan Genomics, Inc. (110053-192 Allegro Targeted Genotyping V2 50-100k; 10053-384 Allegro Targeted Genotyping V2 50-100k). DNA from CD and healthy-derived organoids were sequenced per Tecan’s instructions at the UCSD IGM Genomics Center. The sequencing data was processed by Interval Bio (San Diego, CA), a partner of Tecan Genomics, so that total SNPs and SNPs of special interests could be counted. More specifically, Interval Bio provided the bioinformatics service to list the single nucleotide polymorphism of genotype calls and variant calls with the rsIDs.

##### Treatment of EDMs with probiotic and postbiotic agents

The single cells from non-stricturing non-penetrating (NSNP) patient derived-organoids were seeded in 24-well transwells and allowed to differentiate for 2 days such that the TEER measurement (assessed manually, as described above) reached a stable plateau (indicative of maximal polarization of the monolayer)*. Escherichia coli* Nissle (EcN) was cultured using the same method as *AIEC-* LF82 and added to the apical side of the EDMs with a MOI of 100 and allowed to co-incubate for 8 h. The TEER was measured before and after the treatment at regular time intervals. To determine the impact of postbiotic Hylak Forte (HF), EDMs were preincubated for 12 h with HF and the TEER measurements were carried out for 24 h post-treatment at regular time intervals.

##### Assessment of DNA damage by flow cytometry

DNA double-strand breaks (DSBs) were measured by flow cytometry by detecting *γ*-H2AX in organoids after modifying the published methods ^91^. Briefly, single cell suspensions from healthy and CD colonic organoids were washed using FACS buffer (PBS, 5% FBS, 2 mM Sodium Azide) and fixed/stained using the Biolegend Cyto-Fast Fix Perm buffer set according to the manufacturer’s instructions. Cells were washed in Cyto-Fast Perm wash solution before incubation with the primary antibody anti-g*H2AX* (1:100) for 30 min at room temperature. Cells were washed with Cyto-Fast Perm wash solution, followed by incubation with secondary antibody Alexa Fluor 488 conjugated goat anti-mouse IgG and propidium iodide for 30 min in the dark at room temperature. Cells were washed with Cyto-Fast Perm Wash Solution, resuspended in FACS buffer and data was acquired on a Guava easyCyte flow cytometer (see ^Key^ ^Resource^ ^Table^). Data was analyzed using the FlowJo software (see Key Resource Table).

##### Assessment of cellular senescence using SPIDER β-Galactosidase assay

Senescence associated β-Galactosidase (SA-βGal) was measured in healthy and CD colonic organoids following published protocols^68^. Briefly, organoids were cultured for 7 days and incubated with the 100nM Bafilomycin A1 for 1 hour to inhibit endogenous β-galactosidase activity. Organoids were washed using HBSS (without Ca2+/Mg2+) and incubated with 1 µM of SPIDER-βGal diluted in HBSS for 60 min at 37°C. Organoid were again washed with HBSS (without Ca2+/Mg2+) and disassociated to single cell suspensions with TrypLE and followed by filtering through 70 μm cell strainer and cells were immediately acquired on a Guava easyCyte flow cytometer (see Key Resource Table). Data was analyzed using FlowJo software (see Key Resource Table).

##### Multiplex immunoassays for quantification of cytokines

The cytokines were quantified from the supernatants collected from organoids and EDMs using customized Meso Scale Discovery (MSD)V-PLEX cytokine panels as per the manufacturer’s instructions. All data was obtained using a MESO QuickPlex SQ 120 instrument (see Key Resource Table) and analyzed using MSD® DISCOVERY WORKBENCH 4.0 software (see Key Resource Table).

##### Bacterial clearance assay in CD colonic organoids and the impact of PPAR α/γ dual agonist

Adherent Invasive *Escherichia coli* strain LF82 (AIEC-*LF82*), isolated from the specimens of Crohn’s disease patient, was obtained from Arlette Darfeuille-Michaud^70^. Bacterial clearance assay was performed following our published work^22, 46^. AIEC-*LF82* was grown in LB broth for 8 h under aerobic conditions and then under oxygen-limiting conditions overnight. Healthy and grey cluster EDMs were infected apically with AIEC-*LF82* with a multiplicity of infection (MOI) of 30 for 3 h in presence or absence of 1 µM PPAR α/γ dual agonist (PAR5359) with 16 h preincubation of the drug prior to infection. Gentamicin protection assaywas performed after 3 h with 200 μg/ml of gentamicin for 90 minutes, followed by serial dilution in 1x PBS and plating on LB agar plates. Colonies were counted the next day to measure colony forming unit per ml (cfu/ml).

#### Computational and bioinformatics approach

##### RNA sequencing and data processing

Raw FASTQ files were trimmed, filtered, and mapped to human genome for downstream quantification analyses. Low-quality sequences were trimmed or removed with Trimmomatic. STAR (version 2.6.0a) was used to align reads on the reference genome (human genome hsGRCh38_94). The resulting transcriptome-aligned sequences were used for expression quantification by using RSEM (version 1.3.3) with “–forward-prob 0” option. TPM scores for each sample were used through the RSEM tables (RSEM gene.results tables). We used log2(TPM) if TPM > 1 else TPM - 1 values for each sample as the final gene expression value for downstream expression analyses.

##### RNASeq data analysis

RNASeq data was processed using traditional differential expression analysis tools (DESeq2), clustering techniques (PCA, Hierarchical Agglomerative Clustering), and visualized using heatmaps on R statistical software version 4.1.0 (2021-05-18). Principal component analysis (PCA) was performed in all samples based on bulk RNA seq data using *FactoMineR::PCA* function from R package:*FactoMineR*. Clusters of correlated variables were discovered using Hierarchical Clustering on Principal Components (*FactoMineR::HCPC*) analysis and highlighted using a factorial map (*FactoMineR* v2.5; *factoextra* v1.0.7; *ggplot2* v3.3.5). HCPC dendrograms were visually analyzed to find suitable number of clusters. Heatmaps with dendrograms were generated using python *clustermap* function from the *seaborn* package. The sample clusters description by genes (**Supplementary Figure 3G**) is established using *catdes* function from *FactoMineR* which performs a simple Chi-Square test with significance threshold = 0.05.

##### StepMiner analysis

StepMiner is an algorithm that identifies step-wise transitions using step function in a time-series data^92^. StepMiner undergoes an adaptive regression scheme to verify the best possible up and down steps based on sum-of-square errors. The steps are placed between time points at the sharpest change between expression levels, which gives us the information about timing of the gene expression-switching event. In order to fit a step function, the algorithm evaluates all possible steps for each position and computes the average of the values on both sides of a step for the constant segments. An adaptive regression scheme is used that chooses the step positions that minimize the square error with the fitted data. Finally, a regression test statistic is computed as follows:

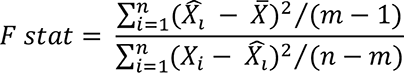

Where 𝑋_𝑖_ for 𝑖 = 1 to 𝑛 are the values, 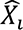 for 𝑖 = 1 to 𝑛 are fitted values. m is the degrees of freedom used for the adaptive regression analysis. 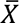 is the average of all the values: 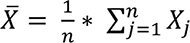. For a step position at k, the fitted values 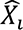 are computed by using 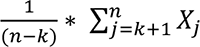 for 𝑖 = 1 to 𝑘 and 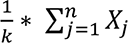 for 𝑖 = 𝑘 + 1 to 𝑛.

##### Composite gene signature analysis using Boolean Network Explorer (BoNE)

Boolean network explorer (BoNE) provides an integrated platform for the construction, visualization and querying of a gene expression signature underlying a disease or a biological process in three steps (**Fig 6, S4**): First, the expression levels of all genes in these datasets were converted to binary values (high or low) using the StepMiner algorithm. Second, Gene expression values were normalized according to a modified Z-score approach centered around *StepMiner* threshold (formula = (expr - SThr)/3*stddev). Third, the normalized expression values for every genes were added together to create the final score for the gene signature. The samples were ordered based on the final signature score. Classification of sample categories using this ordering is measured by ROC-AUC (Receiver Operating Characteristics Area Under The Curve) values. Welch’s Two Sample t-test (unpaired, unequal variance (equal_var=False), and unequal sample size) parameters were used to compare the differential signature score in different sample categories. Violin, Swarm and Bubble plots are created using python seaborn package version 0.10.1. Pathway analysis of gene lists were carried out via the Reactome database and algorithm^80^.

### QUANTIFICATION AND STATISTICAL ANALYSIS

All experiments were repeated at least three times, and results were presented either as one representative experiment (when images are displayed) or as average ± S.E.M (when displayed as graphs). Statistical significance between datasets with three or more experimental groups was determined either using one-way ANOVA including a Tukey’s test for multiple comparisons, or t-tests (Welch’s or Mann-Whitney), as indicated. For all tests, a P-value of 0.05 was used as the cutoff to determine significance and the real p-values are indicated in each figure. For all experiments, the statistical analyses were performed using GraphPad prism 6.1.

For computational analyses, the statistical tests were performed using R version 3.2.3 (2015-12-10). Standard t-tests were performed using python scipy.stats.ttest_ind package (version 0.19.0).

