## Supplemental File for "A Living Organoid Biobank of Crohn’s Disease Patients Reveals Molecular Subtypes for Personalized Therapeutics"

* Equal contribution
‡ Equal contribution

### † Senior corresponding authors:

### [¶]Lead contact author:

### INVENTORY OF SUPPLEMENTARY ONLINE MATERIALS

- Supplementary Text
- Supplementary Figure and Legends (12)
- References Cited

### SUPPLEMENTARY TEXT

#### The two molecular subtypes of CD may display both shared and distinct epithelium-intrinsic defects

The transcriptome analyses revealed a plethora of other clues. Using a previously defined 15-gene crypt-axis score[1] (**Supplementary Figure 4A**) we found the axis in CD-PDOs to be skewed towards differentiation, both in 3D (**Supplementary Figure 4B**-*left*) and 2D (**Supplementary Figure 4C**) growth conditions. Differentiation was more pronounced in the CD-PDOs of the gray cluster, i.e., IDICDs (**Supplementary Figure 4B**-*right*). Consistent with this, *CEACAM7*+ terminally differentiated colonocytes[2] were increased in CD-PDOs, and the IDICD-PDOs uniquely contributes to this pattern (**Supplementary Figure 4D**-*left,* **4E**).

A comprehensive assessment of the various subtypes of *LGR5*+ intestinal stem cells (ISCs) using previously published ISC-signatures[3] revealed that MHCII$^{+}$ *Lgr5*$^{+}$ ISCs are downregulated in both subtypes of CD-PDOs, but more significantly in the yellow cluster PDOs, i.e., S2FCD (**Supplementary Figure 4F-G**). Because MHCII$^{+}$ *Lgr5*$^{+}$ ISCs are non-conventional antigen-presenting cells for CD4$^{+}$ T helper (Th) cells, constituting stem cell-immune cell synapses that balance self-renewal and differentiation in the setting of infection and inflammation[3], their reduction suggests a defect in this homeostatic pathway.

Analysis of the abundance of goblet (*MUC2*) and Paneth cell (*LYZ, REG3A*) transcripts suggested that the proportions of these cells may be altered (**Supplementary Figure 5A-B**). CD-PDOs of both subtypes were deficient in *WFDC2* (**Supplementary Figure 5C**)—an antiprotease molecule that is expressed by goblet cells, inhibits bacterial growth, preserves TJ integrity, prevents bacterial invasion and suppresses mucosal inflammation, and is downregulated in the IBD colon[1]. The TJ-genes (**Supplementary Figure 5D**) and DNA damage genes (**Supplementary Figure 5E**) were reduced in both CD subtypes, but the mitotic checkpoint complex genes were reduced exclusively in the S2FCD-PDOs (yellow cluster; **Supplementary Figure 5F**). The DEGs between healthy and CD-PDOs could successfully classify treatment-responders from non-responders (**Supplementary Figure 5G**) even if the samples were collected prospectively (before treatment; **Supplementary Figure 5H**) and regardless of the modality of treatment (**Supplementary Figure 5H-I**). These results suggest that the CD-PDO-derived intrinsic epithelial processes may have predictive value.

To investigate if the transcriptome of the CD-PDOs reflect the previously known genes/risk alleles implicated in CD, we first defined a list of genes nearest to every genetic variant significantly associated with CD based on GWAS (see *STAR methods*; **Supplementary Figure 5J**). Levels of expression of these set of genes were generally suppressed in CD-PDOs (**Supplementary Figure 5J**-*left*) and such suppression was significant in the IDICD PDOs (**Supplementary Figure 5J**-*right*). A comprehensive analysis revealed that many revealed that many of these genes were uniquely dysregulated in one or the other CD subtype (**Supplementary Figure 5K**). Finally, we found that the DEGs between healthy and CD-PDOs could successfully classify treatment-responders from non-responders (**Supplementary Figure 5G**) even if the samples were collected prospectively (before treatment; **Supplementary Figure 5H**) and regardless of the modality of treatment (**Supplementary Figure 5H-I**). These results suggest that the CD-PDO-derived intrinsic epithelial processes may have prognostic value.

### SUPPLEMENTARY FIGURE AND LEGENDS

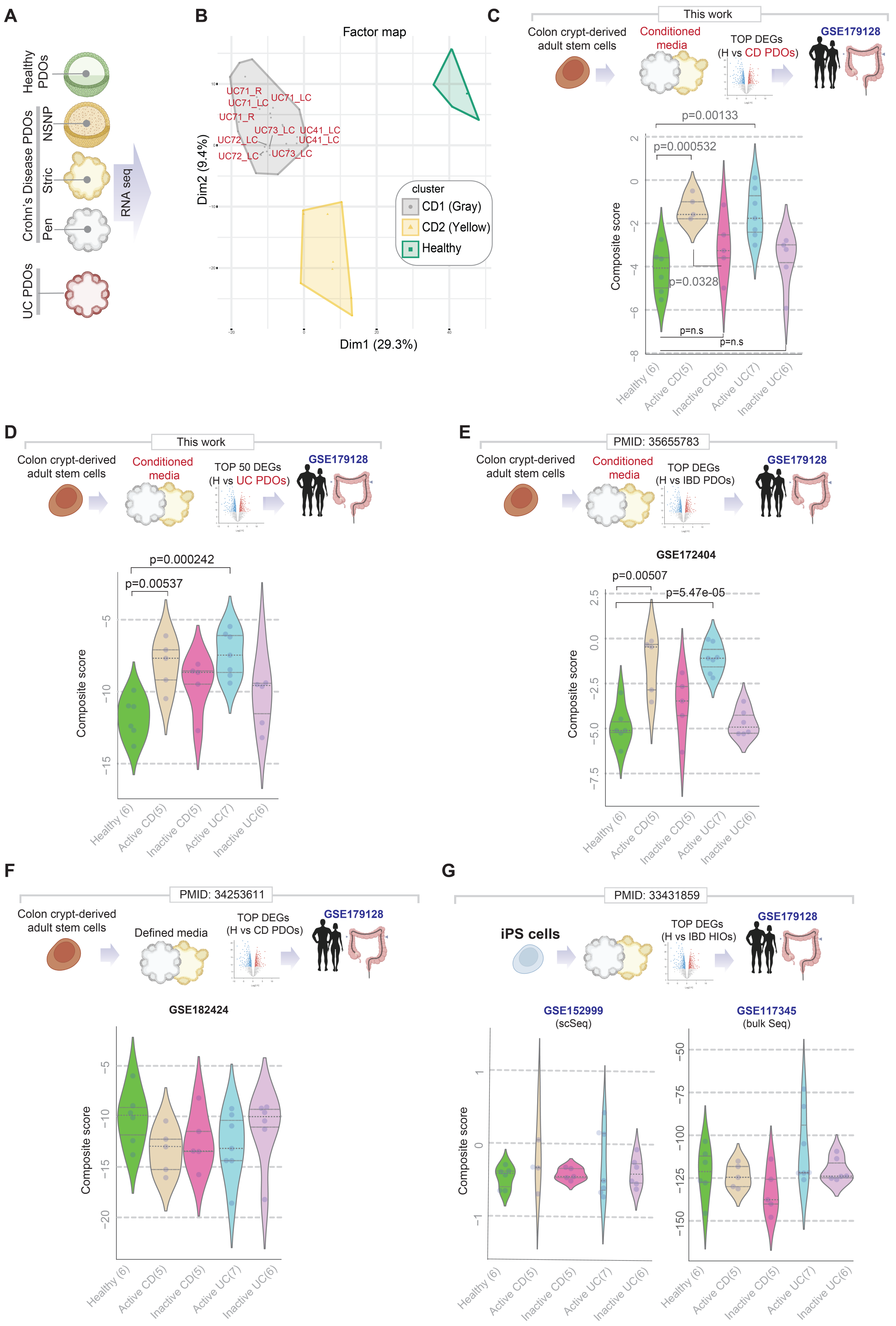


**Supplementary Figure 1 [Related to Figure 2]**

**Benchmarking PDOs as tools for disease modeling: An objective comparison of adult stem cell derived CD-PDOs (this work) against UC-PDOs and other IBD-PDOs from various laboratories.**

**A.** Schematic showing the overall rationale and study design for the transcriptomic studies on healthy and CD-PDOs.

**B**. A factorial map generated by performing the Hierarchical Clustering on principal components (HCPC) analysis is plotted onto the first two dimensions. HCPC was used to compute hierarchical clustering on principal components indicates the division of the H (healthy), UC- and CD-PDOs into three distinct clusters (see **Figure 2B**): gray and yellow, which are differentiated from healthy controls (green). For simplification, CD-PDO annotations are removed, and only UC-PDOs are highlighted. All 10 UC-PDOs analyzed here, were found in the gray cluster with CD-PDOs; none were found in the yellow cluster.

**C**. *Top*: Schematic outlines the strategy used to objectively assess the ability of CD-PDOs to recapitulate the microdissected epithelium from the colons of UC or CD patients, whose disease activities were clinically determined to be active or inactive disease (GSE179128). *Bottom*: Violin plots show the composite score of the PDO-derived top DEGs in the microdissected colonic epithelium. Values in parenthesis indicate unique patients. List of DEGs is provided in **Supplemental Information 4**.

**D.** *Top*: Schematic outlines the strategy used to objectively assess the ability of UC-PDOs to recapitulate the microdissected epithelium from the colons of UC or CD patients, whose disease activities were clinically determined to be active or inactive disease (GSE179128). *Bottom*: Violin plots show the composite score of the PDO-derived top DEGs in the microdissected colonic epithelium. Values in parenthesis indicate unique patients.

**E-G**. *Top*: Schematics outline the strategy used to objectively assess the ability of colonoids, generated either of adult stem cells of healthy or IBD-afflicted colons (E-F) or iPS-cells from healthy or IBD-afflicted patients (G) to recapitulate the microdissected epithelium from the colons of UC and CD patients, whose disease activities were clinically determined to be active or inactive disease (GSE179128). *Bottom*: Violin plots show the composite score of the organoid-derived top upregulated DEGs in each study (indicated with a PMID#) in the microdissected colonic epithelium. Values in parenthesis indicate unique patients.

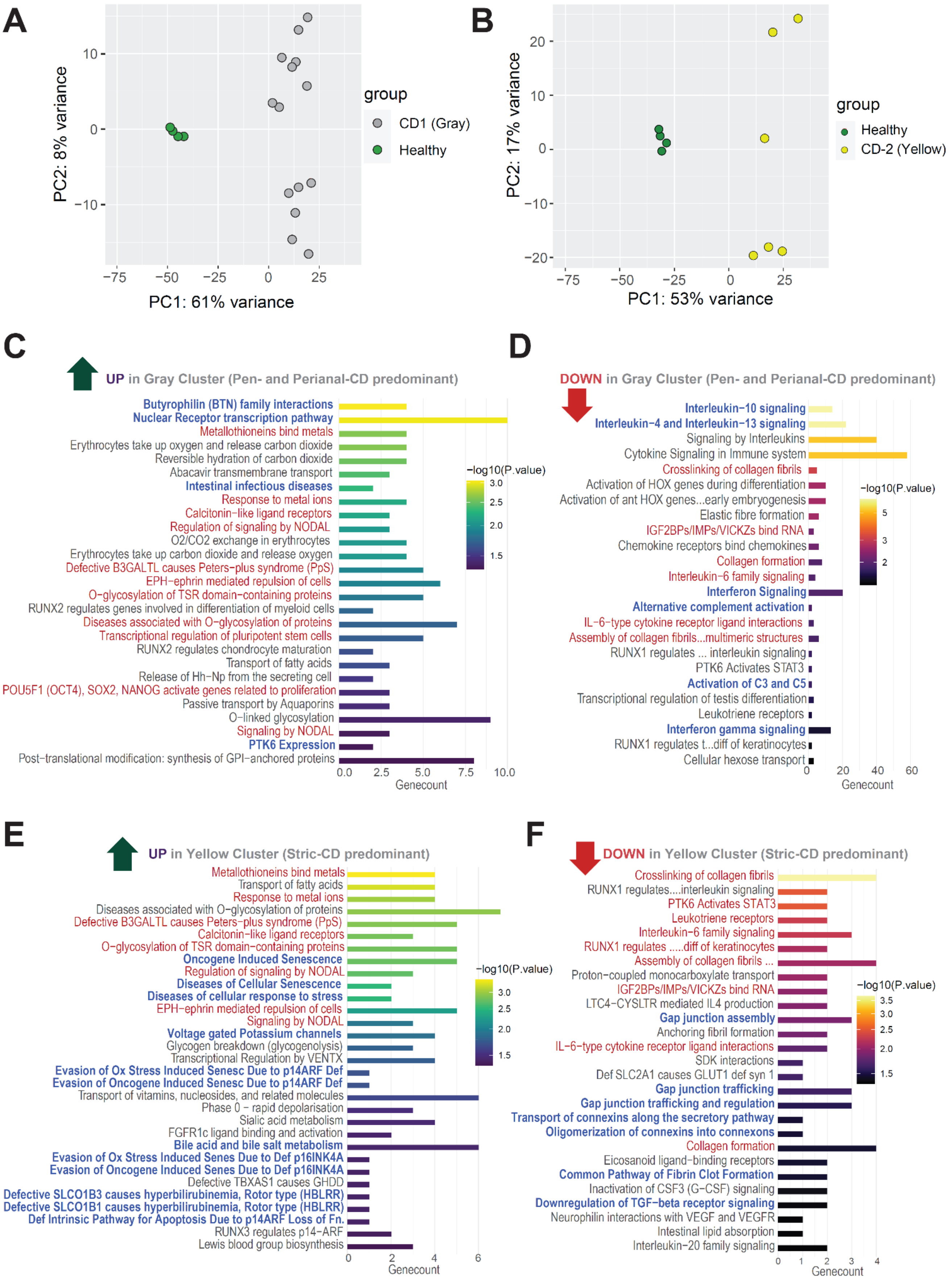


**Supplementary Figure 2 [Related to Figure 2]**

**Analysis of differentially expressed genes (DEGs) and cellular processes and pathways in in the CD-PDOs (gray or yellow clusters) vs healthy PDOs.**

**A-B**. Two-dimensional PCA plots showing clustering of the healthy control PDOs (green) and CD-(molecular) subtypes (gray or yellow) samples.

**C-F**. Enriched pathways from the differential gene expression analyses by DESeq2 with log2 fold changes (LFC)=1 and false discovery rates (FDR) = 0.05. Reactome pathway analysis of Up/Down-regulated genes in the gray/yellow cluster. The vertical axes are the enriched pathways, and the horizontal axes are the number of DE genes in each enriched pathway.

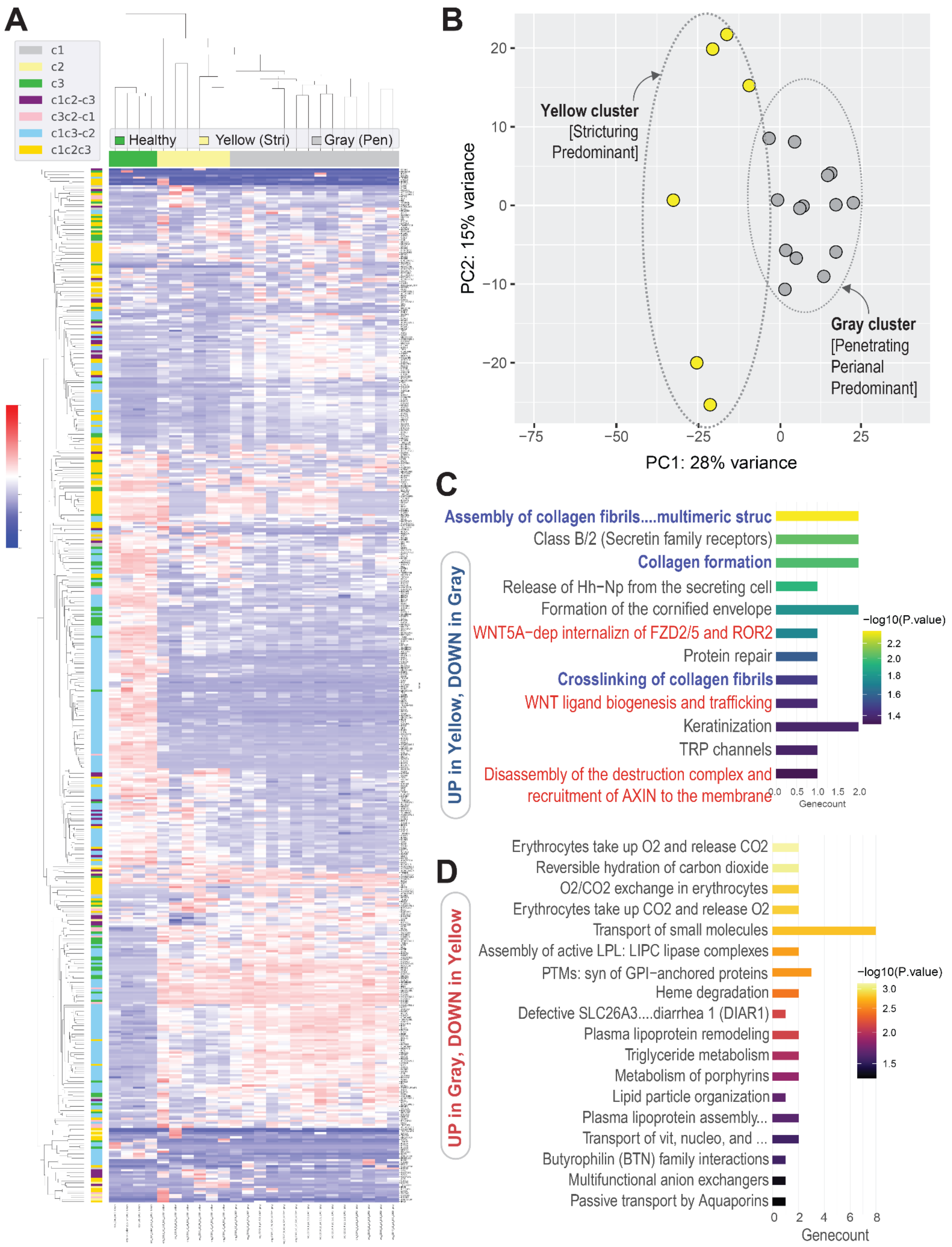


**Supplementary Figure 3 [Related to Figure 2]**

**Analysis of differentially expressed genes (DEGs) and cellular processes and pathways in gray vs yellow CD-PDOs.**

**A**. Unsupervised hierarchical clustering of the top 500 most genes with the highest variance used in clustering by PCA in **Figure 2B**. C1 shows the genes that describe the cluster of gray samples or characterize the CD-PDOs in the gray cluster. C2 shows the genes that describe the CD-PDOs in the yellow cluster. C3 shows the genes that describe the healthy PDOs in the green cluster. C1C2_C3 shows the genes that describe both gray and yellow CD-PDO samples but not the PDOs in the healthy green cluster. C1C2C3 shows the genes that describe all clusters.

**B**. Two-dimensional PCA plots showing clustering of the gray and yellow CD-molecular subtypes

**C-D**. Enriched pathways from the differential gene expression analyses between gray and yellow CD-Subtypes by DESeq2 with log2 fold changes (LFC) =1 and false discovery rate (FDR) = 0.05. Reactome pathway analysis of Up/Down-regulated genes in the gray cluster.

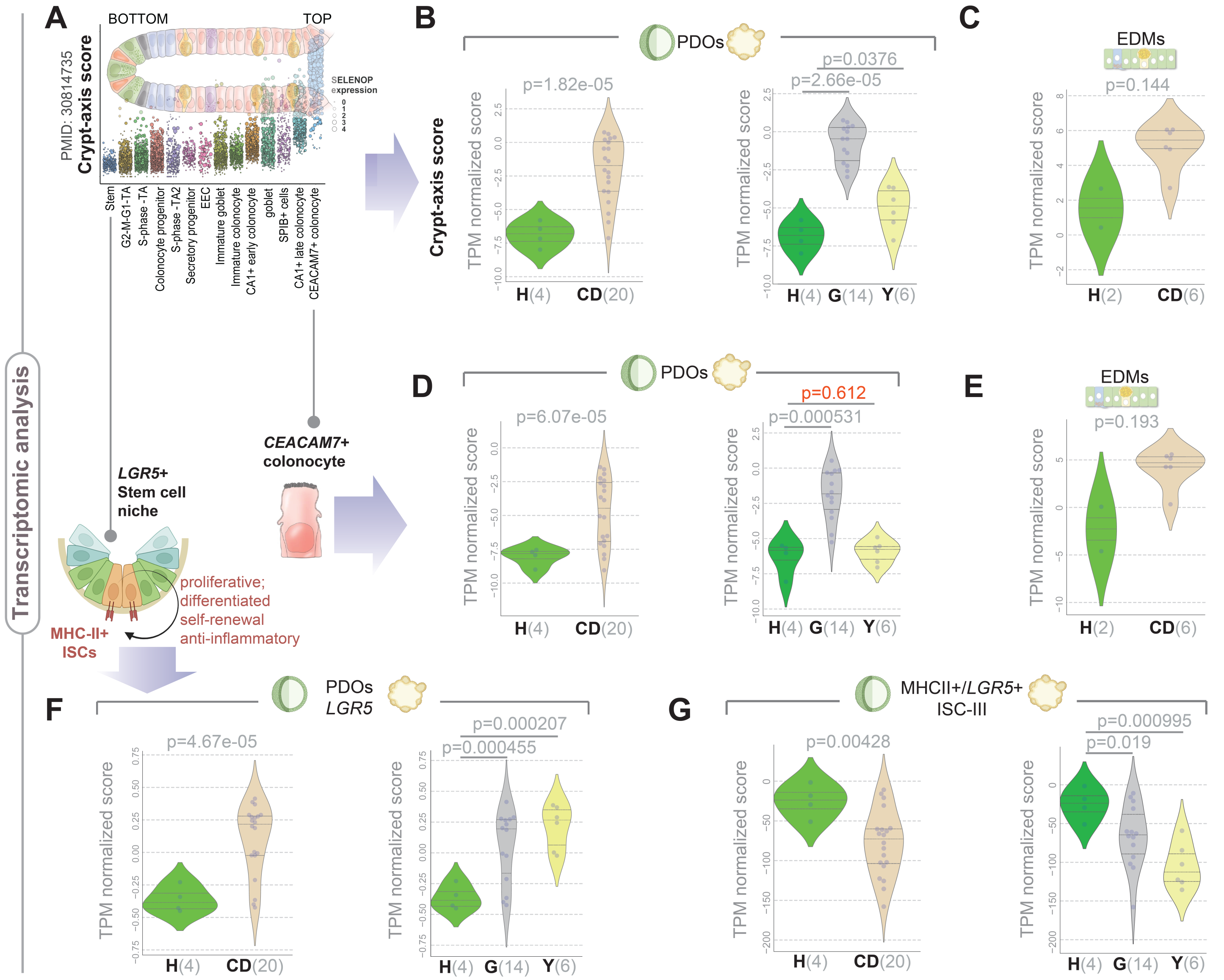


**Supplementary Figure 4 [Related to Figure 2]**

**Two molecular subtypes of CD show shared and unique epithelium-intrinsic defects**.

**A.** Schematic shows the pseudo-spatial distribution of developing epithelial cells along the crypt-villus (base-top) axis. The axis score was derived by using the expression of selected crypt-villus axis markers as defined previously[1,4].

**B-C.** Violin plots show composite score of the set of genes that define crypt-axis score in CD-PDOs, grown either as 3D cultures (B) or differentiated into 2D EDMs (C). Statistical significance was determined by Welch's t-test. H, healthy; G, gray cluster; Y, yellow cluster.

**D-E**. Violin plots show composite score of the set of genes that define terminally differentiated *CEACAM7*+ brush border epithelium in CD-PDOs, grown either as 3D cultures (D) or differentiated into 2D EDMs (E). Statistical significance was determined by Welch's t-test. H, healthy; G, gray cluster; Y, yellow cluster.

**F-G.** Violin plots show *LGR5* expression (F) or the expression of a composite score of genes that define a population of intestinal stem cells (G; ISC-III, *MHCII*+/*LGR5*+) in 3D cultures of CD-PDOs. Statistical significance was determined by Welch's t-test. H, healthy; G, gray cluster; Y, yellow cluster.

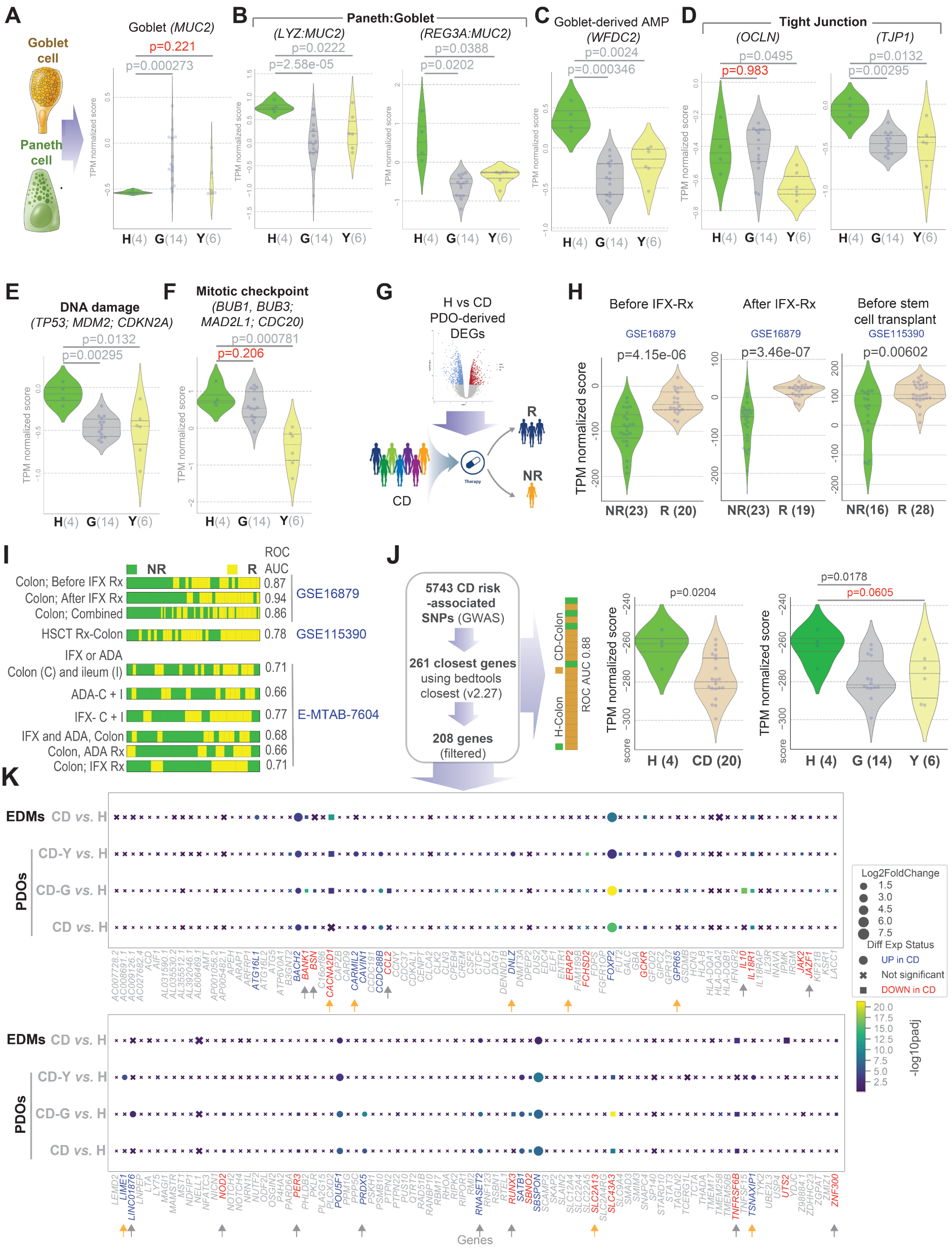


**Supplementary Figure 5 [Related to Figure 2]**

**Integration of transcriptomic information with cellular properties, therapeutic response, and genomics.**

**A-C.** Schematic in A shows the two cell types analyzed in panels A-C. Violin plots show *MUC2* expression (A) or the expression of a composite score of genes representing Paneth and goblet cells (B; *LYZ:MUC2* and *REG3A:MUC2* ratios) and *WFDC2* expression (C) in 3D cultures of CD-PDOs. Statistical significance was determined by Welch's t-test. H, healthy; G, gray cluster; Y, yellow cluster.

**D-F.** Violin plots show the expression of tight junction genes (*OCLN* and *TJP1*) (D), DNA damage response-related genes (E) and genes that function as mitotic checkpoint (F) in 3D cultures of CD-PDOs. Statistical significance was determined by Welch's t-test. Red = insignificant p values. H, healthy; G, gray cluster; Y, yellow cluster.

**G**. Differential expression genes (DEGs) between healthy controls and CD-PDOs were used as gene signature on various publicly available transcriptomic datasets with documented outcome ("responders" vs. "non-responders") of a therapeutic intervention.

**H**. Violin plots show composite score of DEGs in the dataset GSE16879 before (left), after (middle) treatment with the anti-TNFα drug, Infliximab. Violin plots show composite score of DEGs in the GSE115390 dataset (right) where patients with refractory CD received autologous hematopoietic stem cell transplant (HSCT) as the therapeutic modality. R, responder samples; NR, non-responder. Statistical significance was analyzed by Welch's t-test.

**I**. Bar plots show sample order obtained using our signatures (PDO DEGs; see **Supplemental Information 4**) can distinguish responders (R) from non-responders (NR) of therapeutic interventions: (i) before, and after anti-TNFα (GSE16879), (ii) before stem cell transplant (GSE115390), and (iii) with infliximab or adalimumab treatments (E-MTAB-7604). IFX, infliximab; ADA, adalimumab. ROC AUC values are displayed.

**J**. The levels of expression of a list of 208 unique genes nearest to the 5743 CD risk-associated SNPs identified through GWAS [5] that were present also in the current dataset were analyzed. The violin plots show the differences between CD-(molecular)subtypes (gray or yellow cluster). Statistical significances in all panels were determined by Welch's t-test. The gray samples have more of the genes downregulated than the yellow ones.

**K**. Dot plots show the absolute log2 fold change of control samples compared to the CD samples in the four sub-datasets (rows), the differential expression status of CD-associated risk genes (columns) present in all four sub-datasets, and the statistical significance of those genes based on the padj value. The dots' magnitude is absolute (log2 fold change), their shape is the differential expression status, and their colors show the adjusted p-value of the comparison. Blue font = UP in CD over healthy. Red font = DOWN in CD over healthy. Yellow and Gray arrows point to the genes that are uniquely differentially expressed in yellow/gray cluster.

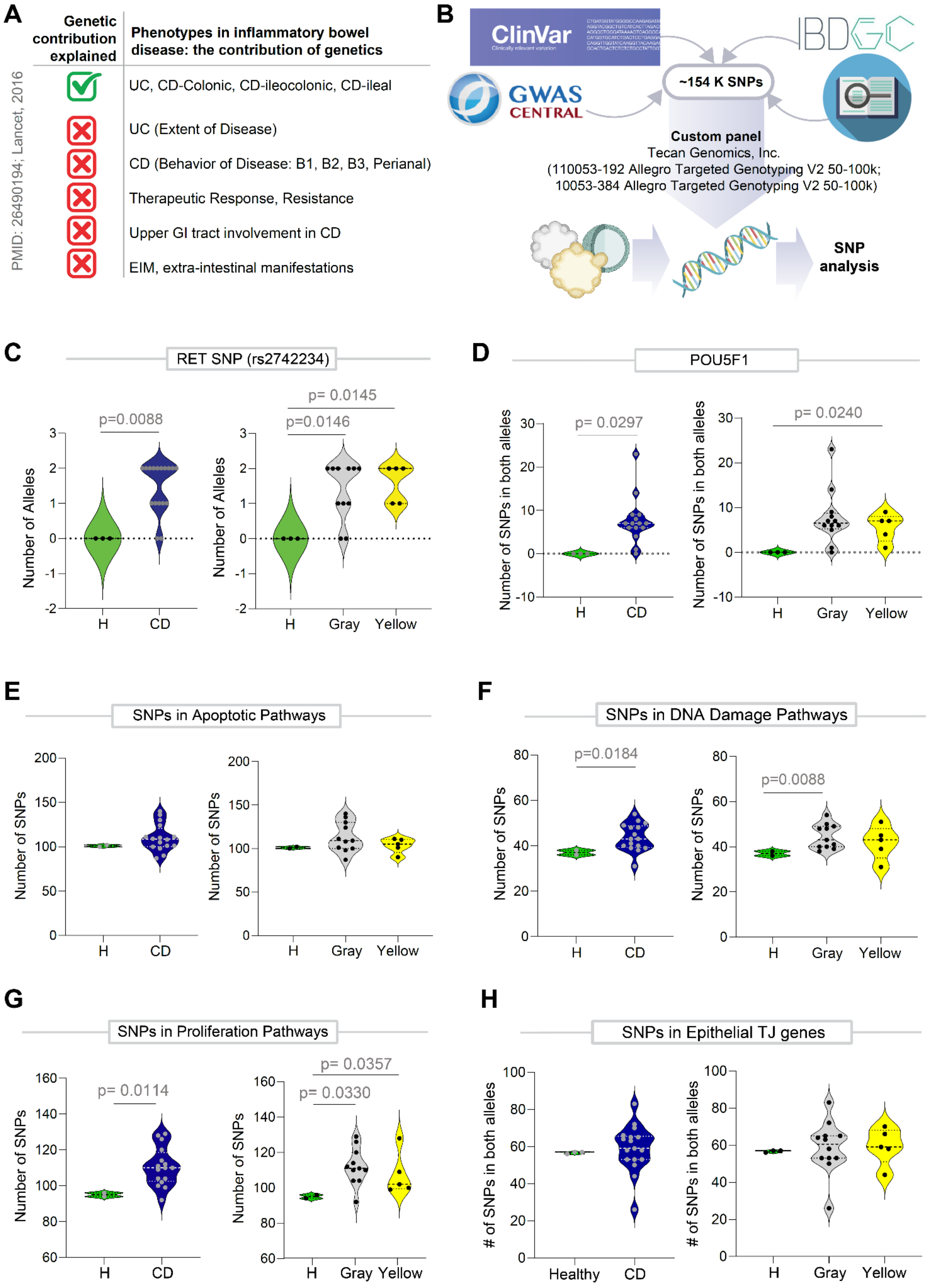


**Supplementary Figure 6 [Related to Figure 2].**

**Genomic analysis of CD-PDOs.**

**A**. Table summarizing the current understanding of the contributions of genetics in CD.

**B**. Schematic showing the key steps in carrying out genomic analyses on healthy and CD-PDOs. See methods for details.

**C-H**. Violin plots display the frequency of SNPs, either in specific genes (C-D; e.g., *RET*, *POU5F1*) or in sets of genes that are involved in various pathways (E-H). Panels on the left compare healthy vs all CD-PDOs, whereas panels on the right compare healthy PDOs against the two molecular subtypes of CD, gray and yellow. Statistical significance was determined by Mann Whitney test. The gene list for SNP analysis is provided as **Supplemental Information 5**.

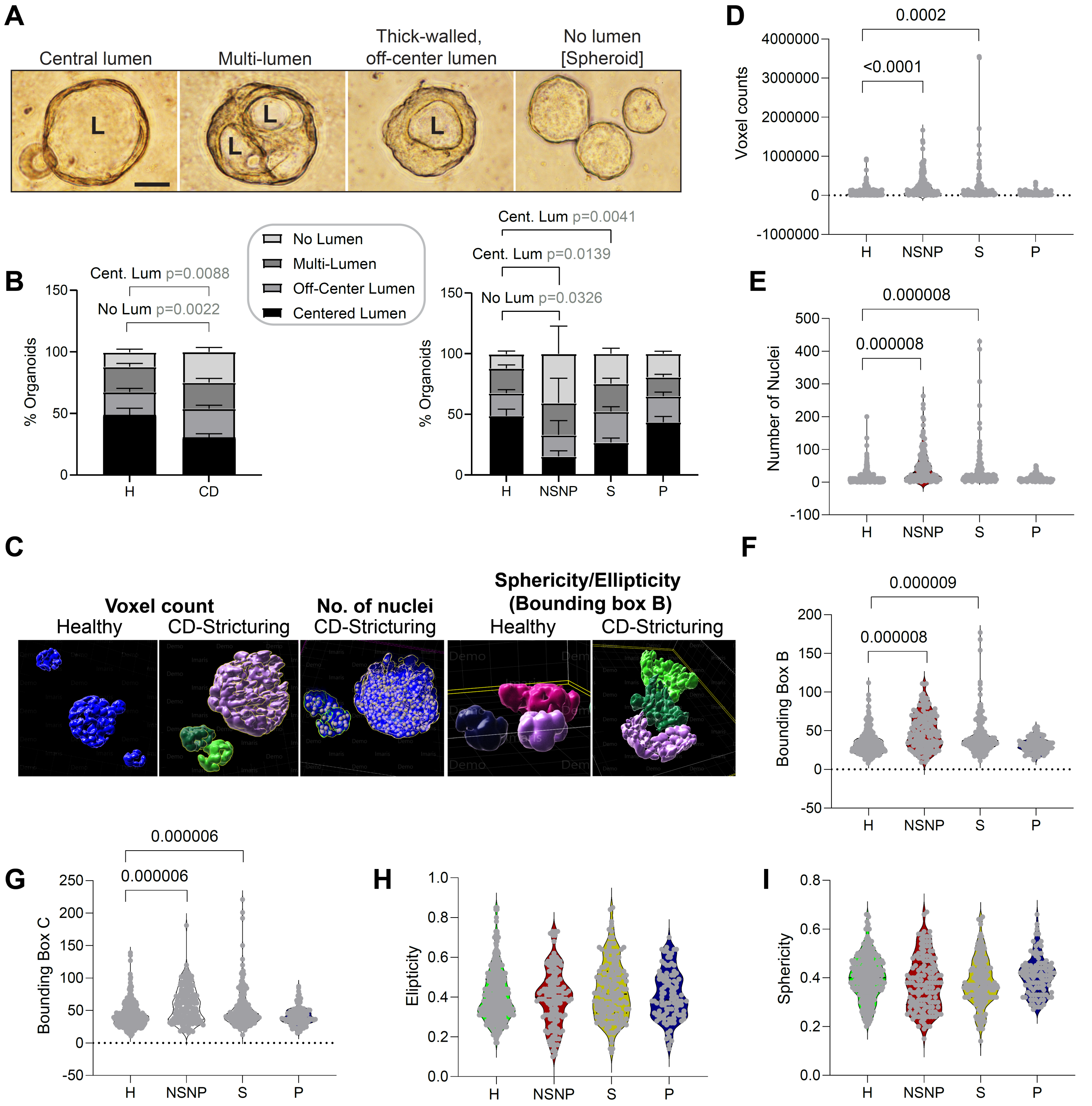


**Supplementary Figure 7 [Related to Figure 3]**

**NSNP (B1)- and stricturing (B2) CD-PDOs show dysmorphic growth.**

**A-B.** Representative images of the 4 major types of organoid structures encountered in 3D cultures of CD-PDOs by light microscopy. Scale bar = 50 µm. L = lumen. Stacked bar plots in B shows the quantification of the proportion of each type of organoid structure in various CD subtypes [B-*left,* all CD subtypes combined; B-*right,* separated into CD subtypes]. Statistical significance was assessed by one way ANOVA. Only significant *p* values are displayed (n = 3-8 in each group).

**C-I.** Quantitative morphometrics were carried out on CD-PDOs using IMARIS. Various parameters were quantified and are represented as violin plots: voxel counts (D), number of nuclei (E), bounding box B (F) and bounding box C (G) and finally, ellipticity (H) and sphericity (I).
See **Supplemental Information 2** for subjects used in each assay.

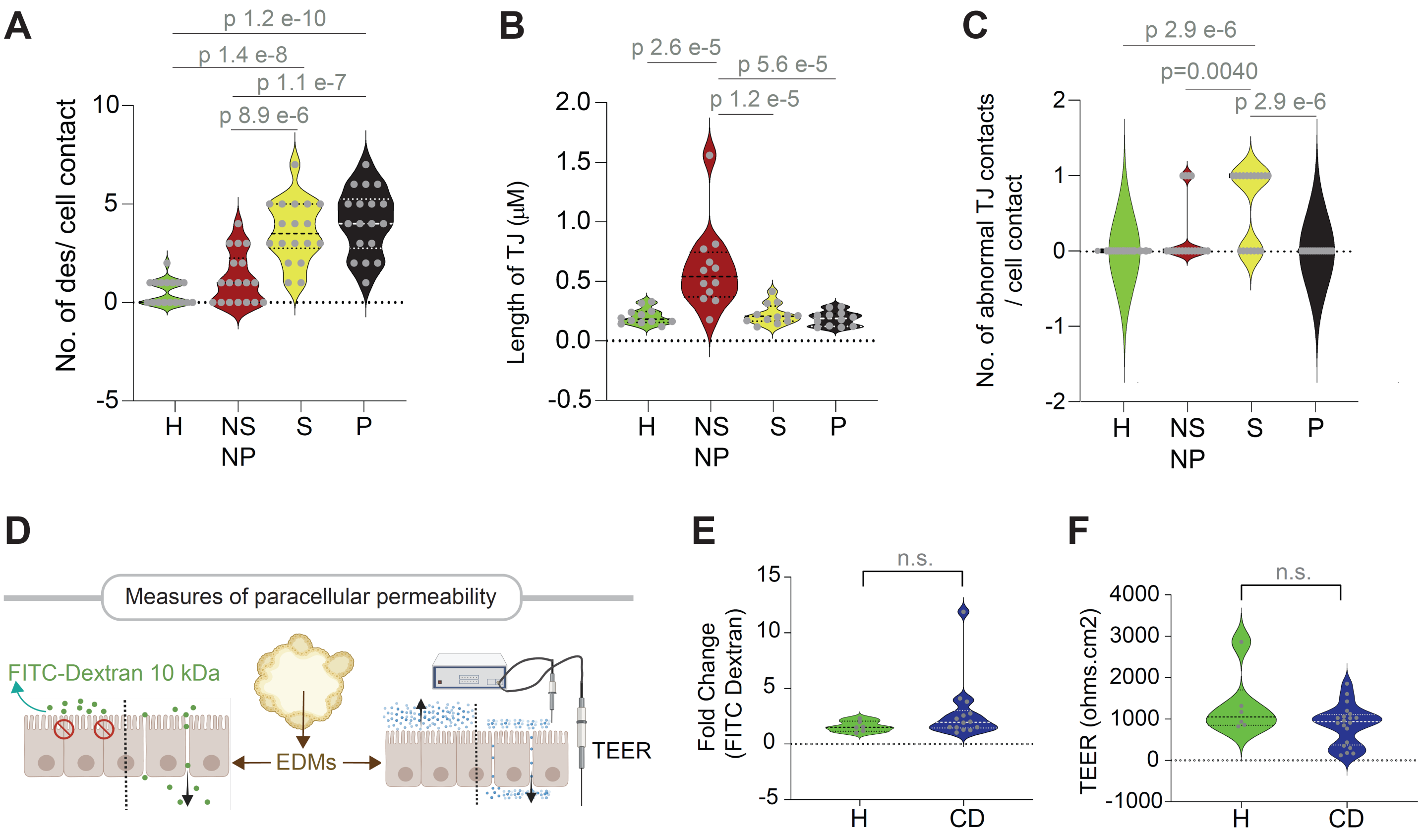


**Supplementary Figure 8 [Related to Figure 3]**

**Characterization of the gut barrier integrity in monolayers derived from CD-PDOs.**

**A-C**. Violin plots display the quantification of no. of desmosomes/cell-cell contact (A), the length of TJ (B) and the frequency of abnormal defects/TJ structure (C) observed by TEM (in **Figure 3C-E**). Statistical significance was assessed by one way ANOVA (n = 7-13 fields analyzed in each subtype of PDO).

**D.** Schematic shows two different approaches used to assess barrier integrity of healthy vs CD EDMs.

**E**. Violin plots show the fold change in FITC dextran leakage in CD-EDMs compared to healthy controls [see **Figure 3J** for each clinical subtype of CD].

**F**. Violin plots show the fold change in TEER in CD-EDMs compared to healthy controls. No statistically significant changes were observed. See **Figure 3** for the visualization of these analyses based on each molecular subtype [**3I-J**] or clinical subtype [**3K-L**] of CD.

See **Supplemental Information 2** for subjects used in each assay.

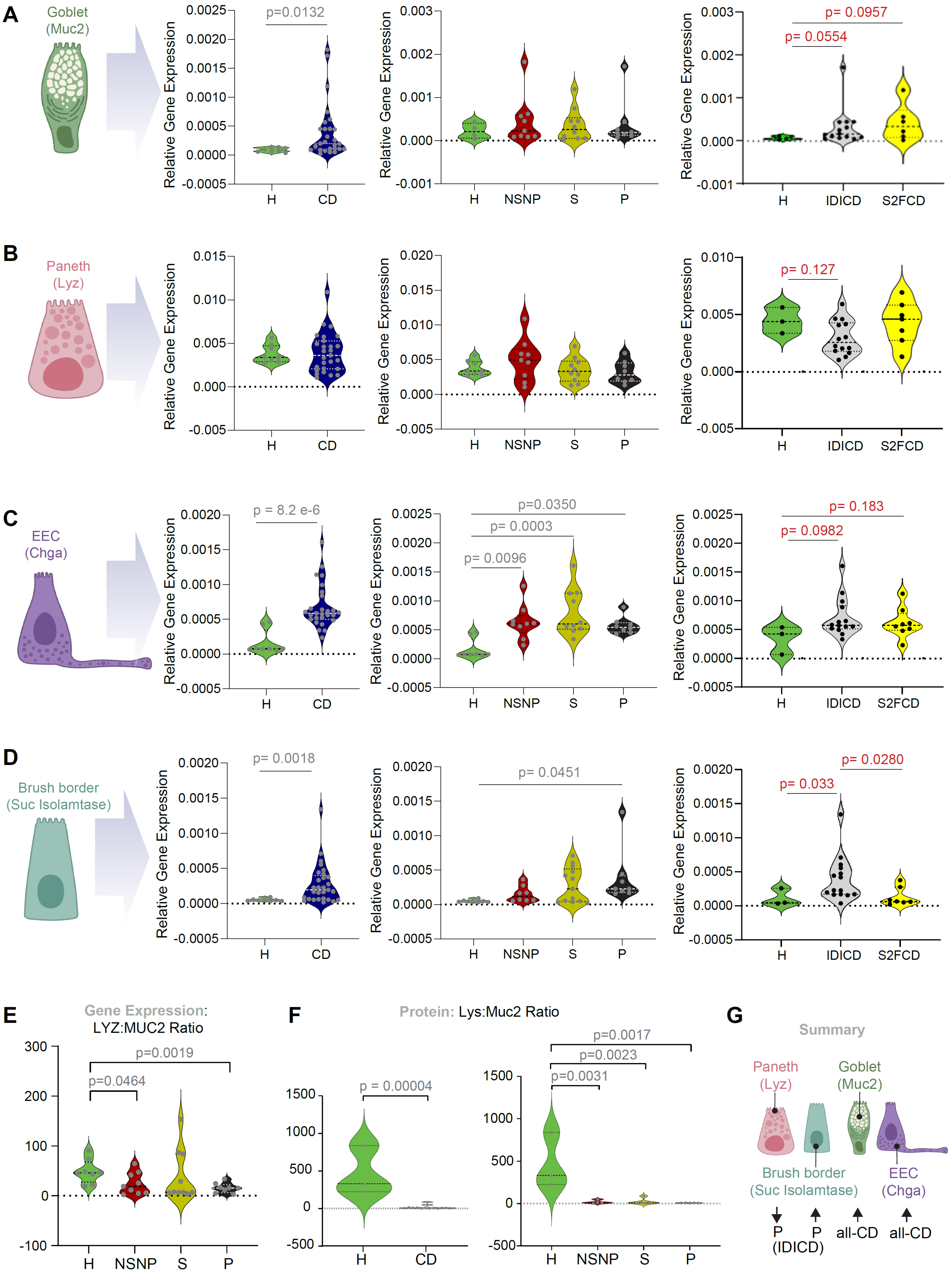


**Supplementary Figure 9 [Related to Figure 4]**

**CD-PDOs retain evidence of altered cell composition.**

**A-D**. Violin plots show the relative abundance of transcripts of *MUC2* (A; a marker of goblet cells), LYZ (B; a marker of Paneth cells), CHGA (C; a marker of enteroendocrine cells) and SI (D; sucrose isomaltase, a marker of brush border cells) in CD-PDOs vs healthy controls [*left,* all CD subtypes combined; *right,* separated into CD subtypes]. Statistical significance was assessed by one way ANOVA [gray font] or Welch's t-test [red font]. Only significant *p* values are displayed (n = 6-15 subjects in each group).

**E**. Violin plots show the ratio of LYZ and MUC2 transcripts (assessed by qPCR) in CD-PDOs vs healthy controls [see **Figure 4B**-*left* all CD subtypes combined; **Figure 4B**-*right,* separated into the two molecular CD subtypes]. Statistical significance was assessed by Mann-Whitney analyses. Only significant *p* values are displayed (n = 6-15 subjects in each group).

**F**. Violin plots show the results of quantification of images in **Figure 4C**, wherein FFPE of CD-PDOs of IDICD subtypes were analyzed for goblet (MUC2; green) and Paneth (Lysozyme; red) cells by confocal immunofluorescence. expressed as ratio of lysozyme to MUC2 in CD-PDOs vs healthy controls [F-*left,* all CD subtypes combined; F-*right,* separated into the clinical subtypes of CD]. Statistical significance was assessed by using unpaired t-test (F-*left*) and one way ANOVA (F-*right*). Only significant *p* values are displayed (n = 3-5 subjects in each group).

**G**. Schematic summarizing the cell type assessment analysis.

See **Supplemental Information 2** for subjects used in each assay.

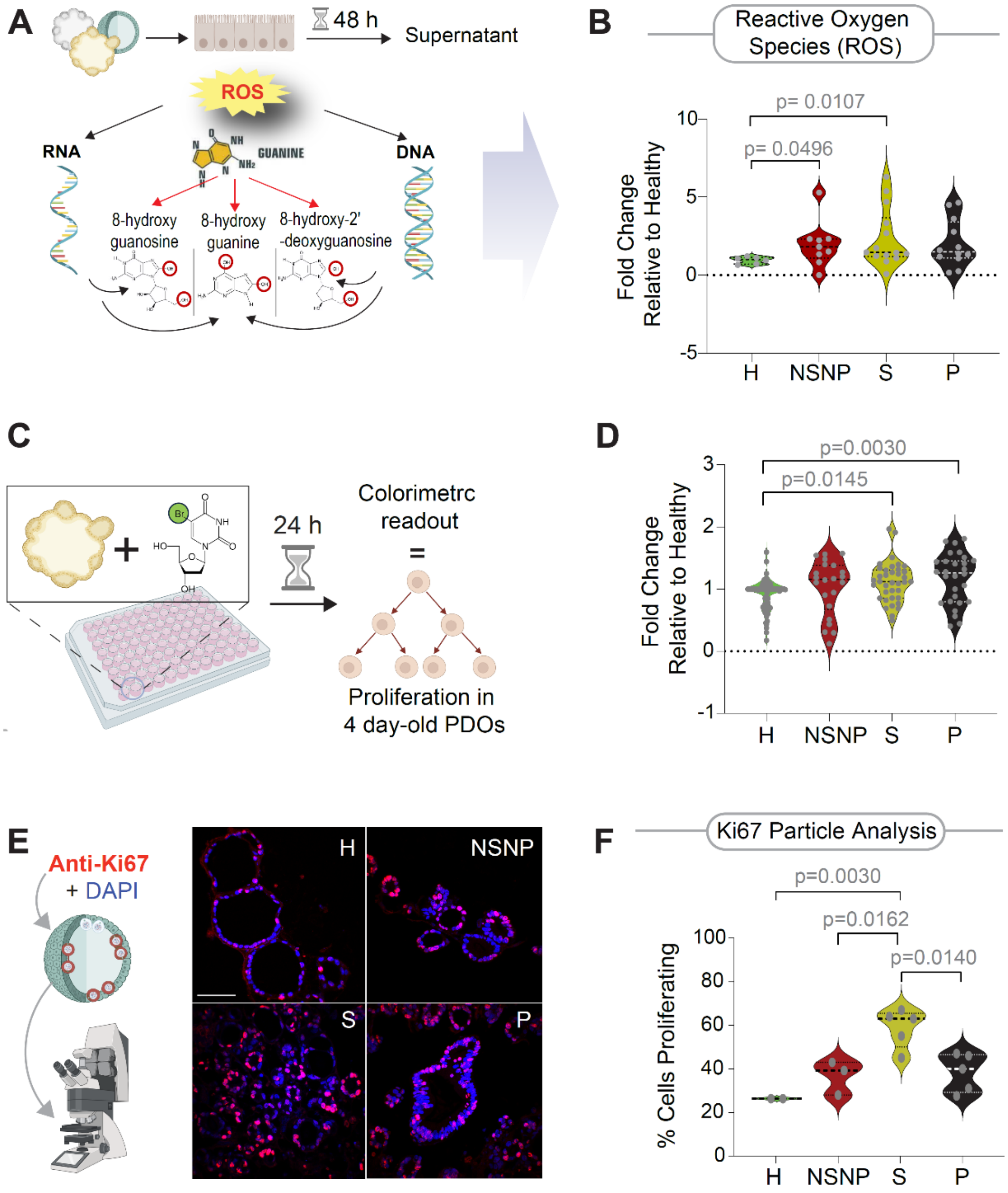


**Supplementary Figure 10 [Related to Figure 4]**

**CD-PDOs retain evidence of high oxidative stress and proliferation.**

**A-B.** Schematic (A) summarizes the assessment of oxidized guanine nucleoside products from damaged DNA/RNA by ELISA. Violin plots (B) show the relative abundance of these products in the 3 clinical subtypes of CD-PDOs vs healthy controls. Statistical significance was assessed by one-way ANOVA. Only significant *p* values are displayed (n = 5-10 subjects in each CD group and n = 3 healthy subjects). See **Figure 4F** for the display of findings as all CD combined or separated into the two molecular subtypes of CD.

**C-D**. Schematic (C) summarizes the proliferation assays performed wherein BrdU-incorporation over 24 h is assessed on four-day old CD-PDOs grown in 96-well plates prior to assessment by ELISA. Violin plots (D) show BrdU incorporation in the 3 clinical subtypes of CD-PDOs vs healthy controls. Statistical significance was assessed by one way ANOVA. Only significant *p* values are displayed (n = 5-8 subjects in each group. H, healthy; NSNP, non-stricturing, non-penetrating; S, stricturing; P, penetrating. See **Figure 4G** for the display of the findings as all CD combined or separated into the two molecular subtypes of CD.

**F-G**. Schematic (F-*left*) summarizes the assessment of Ki67 expression in PDOs by immunofluorescence staining followed by confocal microscopy. Representative images (F-*right*) are shown. Scale bar = 150 µm. H, healthy; NSNP, non-stricturing, non-penetrating; S, stricturing; P, penetrating. Violin plots (G) show % cells with Ki67-positive nuclei in the 3 clinical subtypes of CD-PDOs vs healthy controls, as determined by particle analysis. Statistical significance was assessed by Mann-Whitney

(G-*left*) and one way ANOVA (G-*right*). Only significant *p* values are displayed (n = 2-5 subjects in each group). See **Figure 4H** for the display of the findings as all CD combined or separated into the two molecular subtypes of CD.

It is noteworthy that the penetrating (P, B3) CD-PDOs showed significantly higher BrdU incorporation (**D**), but no significant increase in Ki67 staining (**F**). Because BrdU labels cells during the S phase[6] and is seen in all phases of cell cycle, whereas Ki67 labels cells in all phases except G0[7], findings suggest that penetrating (P, B3) CD-PDOs may have more cells in G0 phase.

See **Supplemental Information 2** for subjects used in each assay.

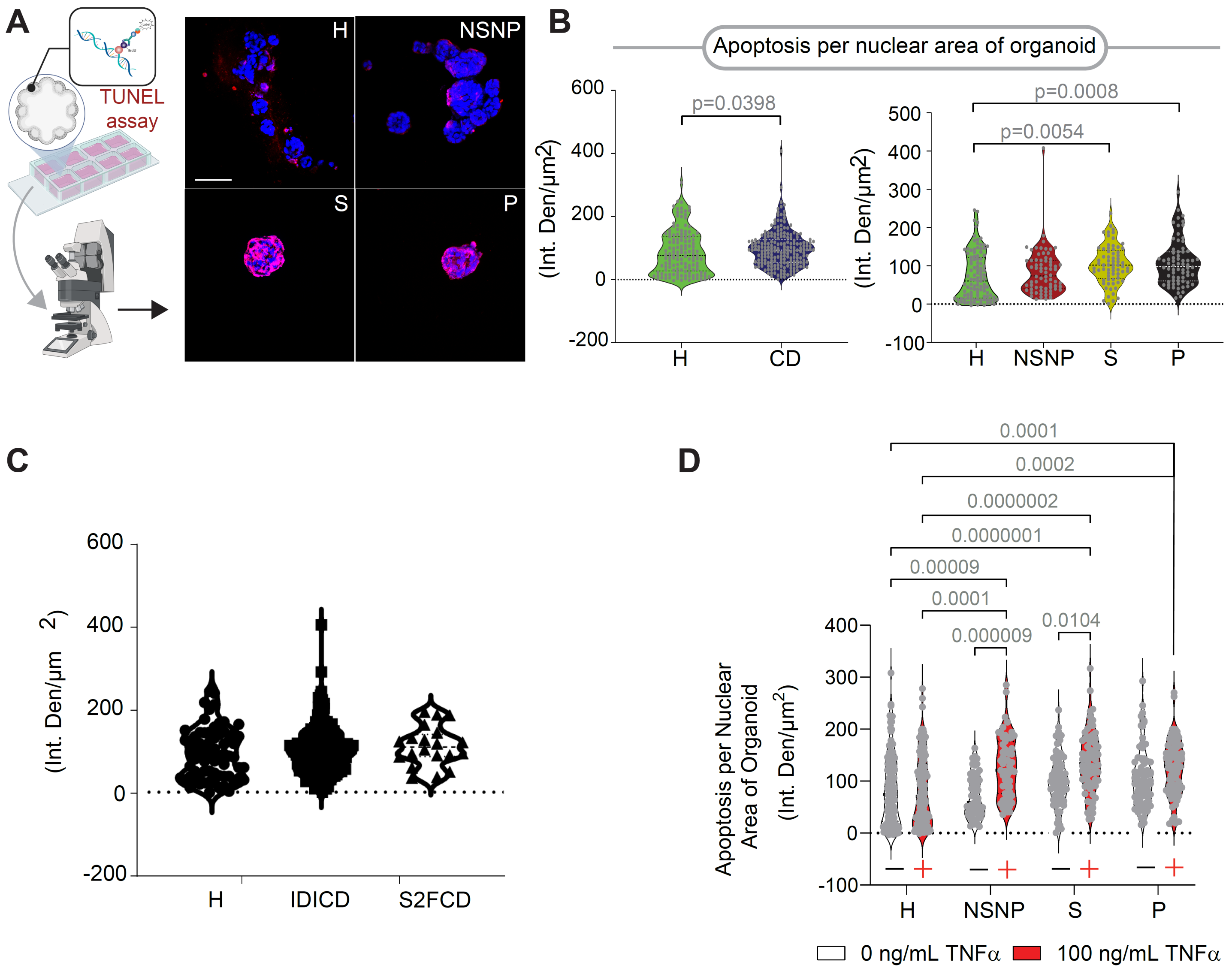


**Supplementary Figure 11 [Related to Figure 4]**

**CD-PDOs retain evidence of high apoptosis.**

**A.** Schematic (A-*left*) summarizes the TUNEL assays performed wherein four-day old CD-PDOs prior to fixation and staining with anti-BrdU (red) and DAPI (nuclei; blue) and analysis by confocal imaging. Representative images (A-*right*) are shown. Scale bar = 150 µm.

**B-C**. Violin plots show densitometry analysis of BrdU incorporation (red pixels) in the nuclei [B-*left,* all CD subtypes combined; B-*right,* separated into the clinical subtypes of CD; C, separated into the molecular subtypes of CD]. Statistical significance was assessed by Mann-Whitney (C-left) and one way ANOVA (C-*right*). Only significant *p* values are displayed (n = 5-6 subjects in each group).

**D**. Findings of TUNEL assays performed wherein four-day old CD-PDOs are challenged with recombinant TNFα for 16 h prior to fixation and staining with anti-BrdU and analysis by confocal imaging. Violin plots show densitometry analysis of BrdU incorporation (red pixels) in the nuclei. Statistical significance was assessed by one way ANOVA (n = 5-6 subjects in each group).

See **Supplemental Information 2** for subjects used in each assay.

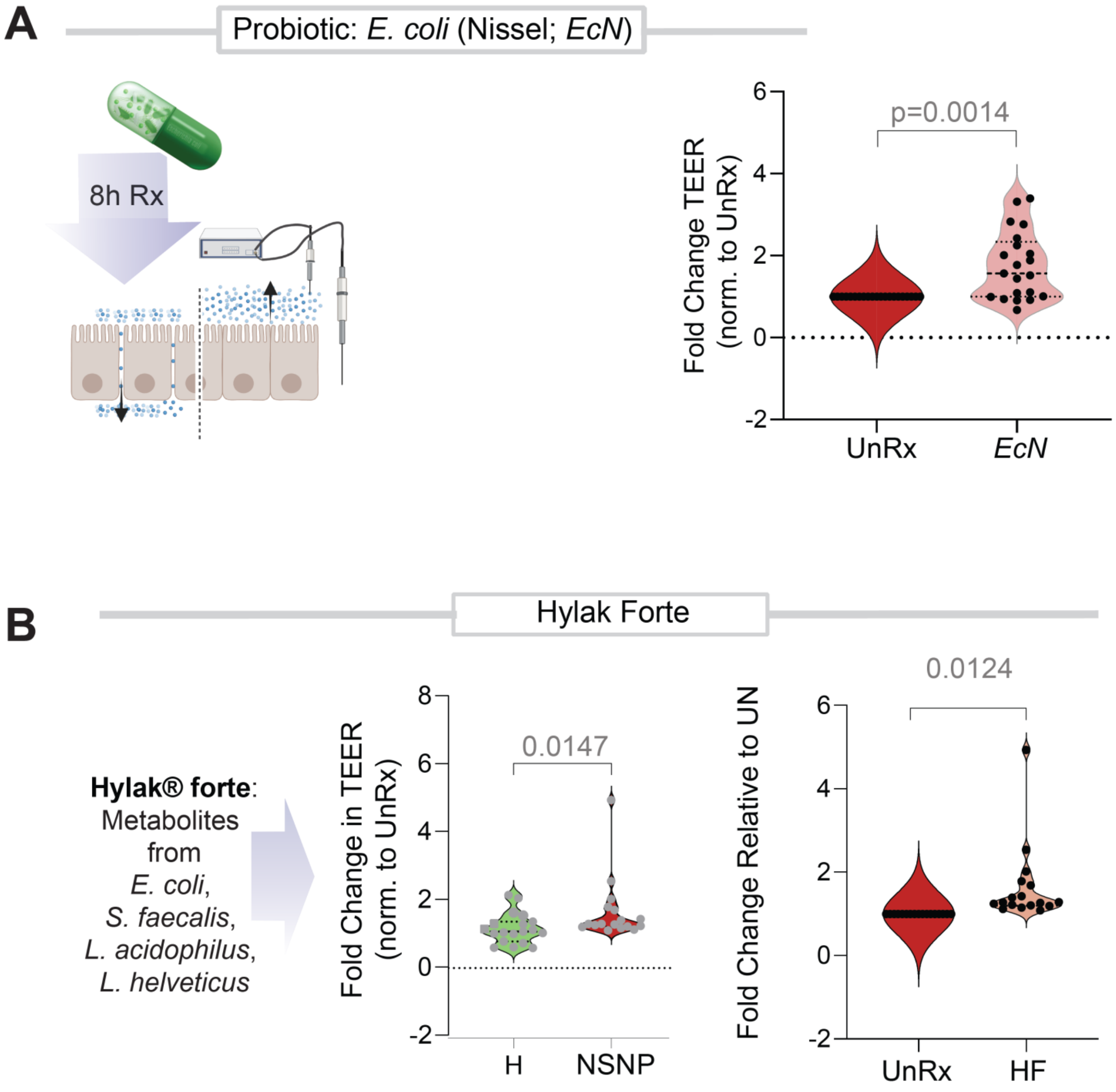


**Supplementary Figure 12 [Related to Figure 5].**

**Barrier defect in NSNP CD-PDOs can be repaired using prebiotics and postbiotics.**

**A-B.** Reversal of the defects in the integrity of the gut barrier observed in NSNP-CD using either prebiotics (A; *E. coli* Nissel, EcN) or postbiotics (B; Hylak Forte®). Violin plots display the fold change in TEER compared to untreated control EDMs. Statistical analysis was performed using paired student t-test.

5. Matt Kanke MMK, Sean Connelly, Matthew Schaner, Michael T. Shanahan, Elisabeth A. Wolber, Caroline Beasley, Grace Lian, Animesh Jain, Millie D. Long, Edward L. Barnes, Hans H. Herfarth, Kim L. Isaacs, Jonathan J. Hansen, Muneera Kapadia6, José Gaston Guillem6, Terrence S. Furey, Shehzad Z. Sheikh, and Praveen Sethupathy. Single-cell analysis of colonic epithelium reveals unexpected shifts in cellular composition and molecular phenotype in treatment-naïve adult Crohn's disease BioRxiv 2021.

6. Matatall KA, Kadmon CS, King KY. Detecting Hematopoietic Stem Cell Proliferation Using BrdU Incorporation. Methods Mol Biol 2018;1686:91-103.

7. Li LT, Jiang G, Chen Q, Zheng JN. Ki67 is a promising molecular target in the diagnosis of cancer (review). Mol Med Rep 2015;11:1566-72.
